# The megabase-scale crossover landscape is independent of sequence divergence

**DOI:** 10.1101/2022.01.10.474936

**Authors:** Qichao Lian, Victor Solier, Birgit Walkemeier, Bruno Huettel, Korbinian Schneeberger, Raphael Mercier

## Abstract

Meiotic recombination frequency varies along chromosomes and strongly correlates with sequence divergence. However, the causality underlying this correlation is unclear. To untangle the relationship between recombination landscapes and polymorphisms, we characterized the genome-wide recombination landscape in the absence of polymorphisms, using *Arabidopsis thaliana* homozygous inbred lines in which a few hundred genetic markers were introduced through mutagenesis. We found that megabase-scale recombination landscapes in inbred lines are strikingly similar to the recombination landscapes in hybrids, with the sole exception of heterozygous large rearrangements where recombination is prevented locally. In addition, we found that the megabase-scale recombination landscape can be accurately predicted by chromatin features. Our results show that polymorphisms are not causal for the shape of the megabase-scale recombination landscape, rather, favor alternative models in which recombination and chromatin shape sequence divergence across the genome.

## Introduction

Meiotic recombination is initiated by the formation of numerous DNA double-strand breaks, a minority of which are repaired as crossovers (COs), resulting in reshuffling of the genetic material between generations, and is thus crucial for diversity, adaptation, evolution, and breeding^1–4^. Two pathways have been described for CO formation (class I and II)^1,3,4^. Class I COs represent the vast majority of COs and are subject to interference, the propensity of COs to be widely spaced along chromosomes^5^.

COs are not homogeneously distributed and recombination frequencies vary along chromosomes^6,7^. Many different features are correlated with the recombination landscape. One consistent pattern across monocentric species is the suppression of COs at and next to centromeres^3,8,9^. The landscape can also differ between the two sexes of the same species, a phenomenon called heterochiasmy^10–13^. Polymorphism between homologs can negatively affect crossovers, as observed very locally at crossover hotspots or even completely suppress crossovers in cases of large polymorphisms, like megabase-scale inversions^7,14–21^. In contrast, however, heterozygous regions in *Arabidopsis thaliana* showed increased recombination rates when juxtaposed with homozygous regions suggesting that the density of small-scale sequence divergence can increase recombination rates^22^. In addition, increasing single nucleotide polymorphism (SNP) density in hybrids associates positively with COs, and the pericentromeric regions that are dense in polymorphisms are also elevated in COs, potentially due to a positive feedback of mismatch recognition during CO formation^21^. A positive correlation between polymorphisms and recombination landscapes can be also observed in natural populations: in many species, historical recombination landscapes as deduced from linkage disequilibrium are positively correlated with SNP densities^23–27^.

To better understand the relationship between polymorphisms and meiotic recombination, we here compared CO distribution along chromosomes in the absence (inbred lines) and presence (hybrids) of polymorphisms. In hybrids, the numerous DNA polymorphisms can be used to precisely map COs^6,28–37^, while this is not an option in homozygous inbred lines. Instead, CO frequency could be estimated by cytological techniques, but this poses several challenges, such as the difficulty in identifying individual chromosomes, low resolution, and the limited number of cells that can be analysed. Alternatively, fluorescence-tagged lines (FTLs) could be used to measure recombination in intervals flanked by markers conferring fluorescence in seeds or pollen grains but are not suitable for mapping the genome-wide CO landscape^38–40^.

In this study, we developed a method to analyse genome-wide recombination landscapes in inbred lines based on the introduction of a few markers by chemical mutagenesis. We characterized the crossover landscapes of two Arabidopsis inbred lines (Columbia (Col) and Landsberg (L*er*)) and compared them to the CO landscape of the Col/L*er* F1 hybrid as well as with the historical recombination pattern in this species. The CO landscapes of the inbred lines were remarkably similar to each other as well as to the historical recombination landscape and that of the hybrid with the exception of local suppression due to large heterozygous rearrangements. This shows that polymorphism density, with the exception of large structural variations, is not a major determinant of the CO landscape.

It has been shown before that COs co-localize with gene promoters and with regions of open chromatin and low levels of DNA methylation^3,30,41,42^. We now show that chromatin accessibility, gene density, and DNA methylation are sufficient to explain 85% of the megabase-scale recombination landscape in Arabidopsis.

## Results

To investigate the landscape of meiotic recombination in *Arabidopsis thaliana* inbred lines, we applied moderate EMS mutagenesis to introduce genetic markers into the genomes of *A. thaliana* Col-0 and L*er*. Independent M_2_ mutants were crossed to generate F1*s, and independent F1*s were reciprocally crossed to generate F1 populations, which were used to analyse recombination independently in female and male meiosis. (Figure 1, Supplemental Figure S1, Supplemental Table S1, Materials and methods). Through Illumina short-read genome sequencing of F1*s and F1s, we identified 838–955 high-confidence mutations segregating in the Col populations and 471–539 in the L*er* populations (Supplemental Table S1), which is a negligible level compared to the natural divergence between these two accessions^14,37,43^. The markers were pretty uniformly distributed across the chromosomes, which allowed the identification of meiotic CO events (Supplemental Figure S2). We analysed four independent pairs of populations of both accessions, with a total of 309 and 309 progeny derived from male and female meiosis in Col and 253 and 251 progenies derived from male and female meiosis in L*er*. Overall, we identified 3,155 COs in Col (examples shown in Supplemental Figure S3 and S4; median resolution 522 kb; Supplemental Table S3) and 2,004 (median resolution 855 kb, Supplemental Table S4) COs in L*er*. We observed a consistent CO frequency between the replicate populations (Figure 2A and 2B), arguing against the unlikely possibility that an EMS mutation dominantly affected CO numbers in the F1*. CO numbers were correlated neither with sequence depth nor with the number of markers, suggesting an absence of bias in CO detection (Supplemental Figure S5). In Col male meiotic cells, CO number was previously estimated to be ∼10.5 with MLH1-HEI10 foci counts^44,45^. However, MLH1 and HEI10 are exclusive markers of class I COs, and do not mark class II COs, which are estimated to represent ∼15% of the total COs^1,46^, suggesting a total number of ∼12.1 COs per male meiocytes. Here, we detected an average of 6.13 COs per Col male gamete, which corresponds to 12.26 COs per male meiocyte, fitting very well with the expected number. Altogether, this shows that our method robustly detects COs in inbred lines.

**Figure 1.**
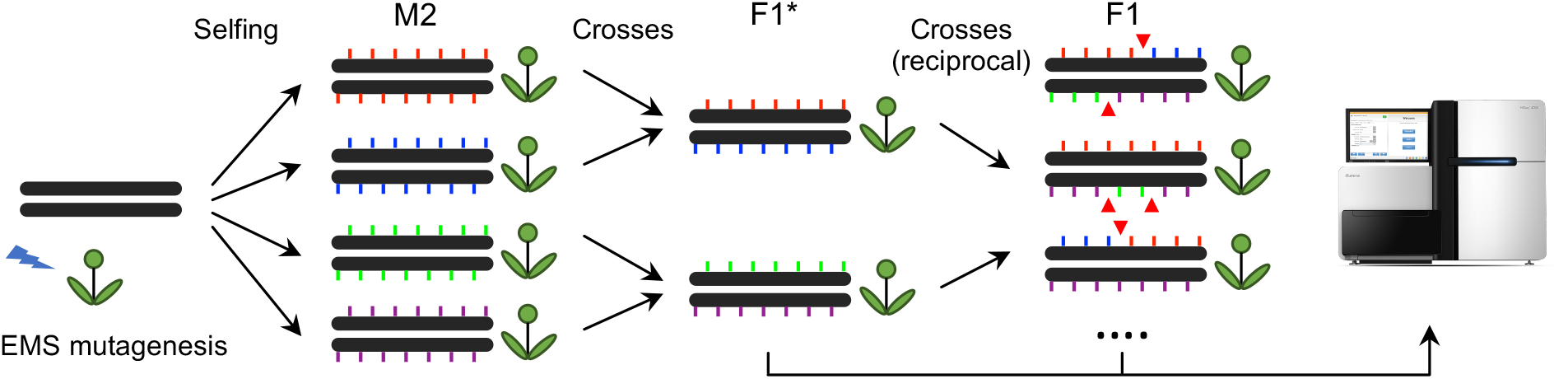
Experimental design for CO identification in Arabidopsis inbred lines. M2 plants were derived from the selfing of independent EMS-treated seeds. Pairs of M2s were crossed to produce F1*s, which are then heterozygous for a set of unique mutations defining two phases, indicated by coloured ticks. Two F1*s were then reciprocally crossed to generate F1 populations. The DNA of leaves of F1* and F1 plants were sequenced using Illumina. The color-coded ticks indicate EMS-induced mutation markers. The red triangles represent COs, detected by phase switching. This design allows the detection of COs that occurred in the male and female meiocytes of the F1* plants.

**Figure 2.**
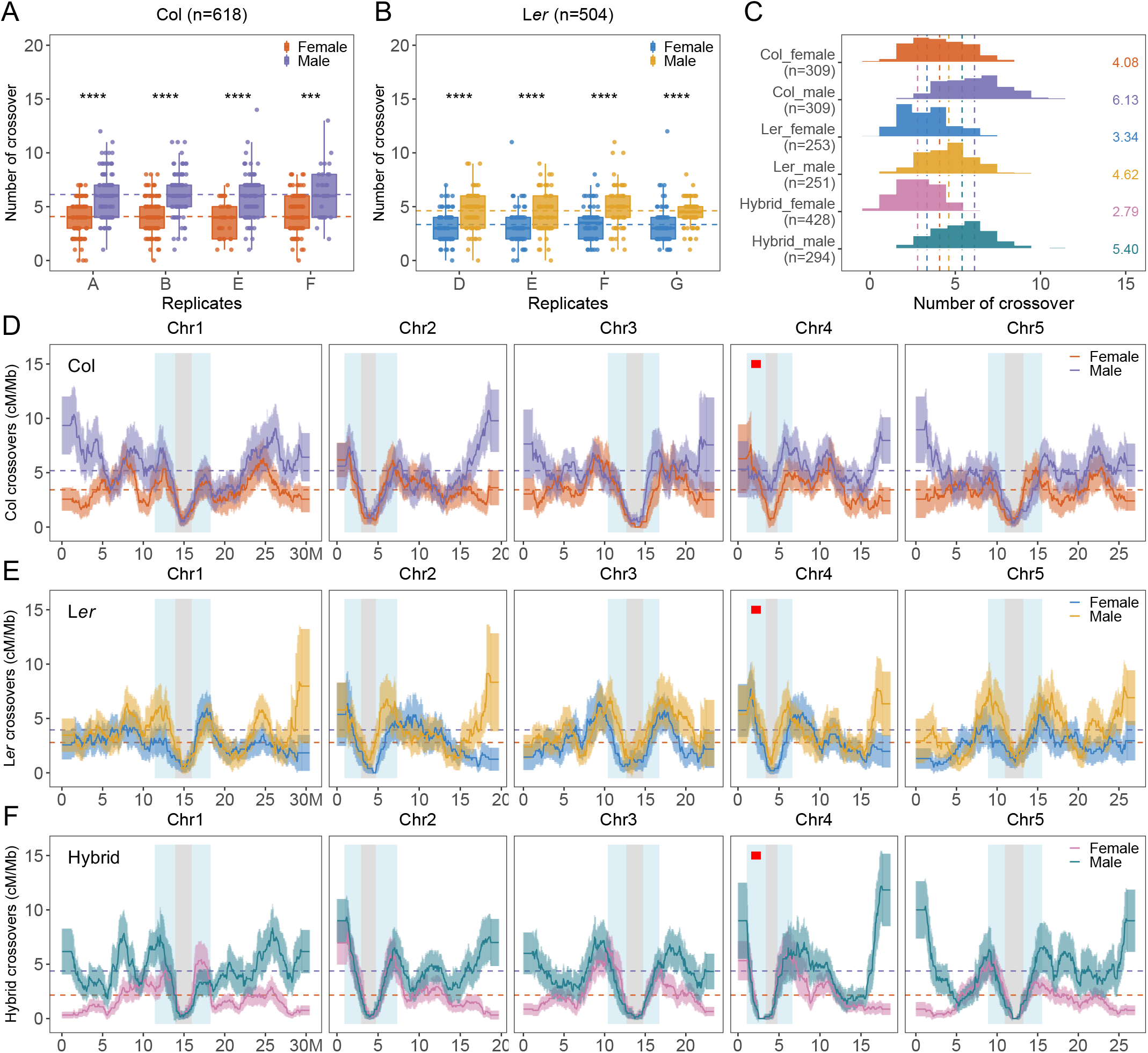
Analysis of female and male COs in Col, L*er*, and F1 hybrid populations. (A–B) The number of COs per gamete in each replicate population of Col and L*er*, respectively. Each letter corresponds to one F1* plant (Figure 1). The Mann-Whitney test was used to evaluate the differences in CO numbers between female and male meiosis. (C) Distribution of CO number per gamete in female and male meiosis of Col, L*er*, and F1 hybrids. The mean number of COs are color-coded and indicated by dashed lines. The sample sizes are indicated in parentheses. (D–F) The chromosomal distribution (sliding window-based, window size 2 Mb, step size 50 kb) of COs in female and male meiosis of Col, L*er*, and F1 hybrids, with 90% confidence interval. The genome-wide mean level of recombination is shown with dashed lines. The pericentromeric and centromeric regions are indicated by grey and blue shading separately. The ∼1.2 Mb inversion between Col and L*er* on chromosome 4 is indicated by a red bar.

To compare the recombination landscapes of Col and L*er* with the corresponding F1 hybrid, we sequenced reciprocal back-crosses of Col/L*er* F1 hybrids with Col to identify COs in 428 and 294 progenies derived from female and male meiosis, respectively. We identified 1,192 COs (median resolution 739 bp) and 1,587 COs (median resolution 1,019 bp) in female and male hybrids, respectively (Supplemental Figure S6, Supplemental Table S5). The female and male high-resolution CO distribution that we obtained is consistent with a previous dataset that described female/male CO landscapes with lower resolution^11^ and CO distribution in the same hybrid in F2s that does not distinguish female and male COs^7^ (Supplemental Figure S7–8, Supplemental Table S6). Comparison of the genomic compartments where CO occurred did not reveal differences between females and males, with COs notably enriched in promotor regions in both sexes. This suggests that the factors driving fine-scale CO placement are similar in female and male meiosis (Supplemental Figure S7D–E).

In all three types of populations, Col, L*er*, and hybrid, we observed heterochiasmy, i.e., significantly more COs in male than female meiosis (Mann–Whitney test, p < 2.2e-16, Figure 2C). In male meiosis, both the highest (Col, 6.13) and the lowest (L*er*, 4.62) numbers of COs are observed in inbred lines, with the hybrid exhibiting an intermediate number of COs (5.4), suggesting that CO number in males is mainly genetically controlled in trans and not by sequence polymorphism. In females, the highest CO number is also observed in Col (4.08), with less COs in L*er* (3.34, p = 1.2e-07), indicating that the same trans mechanism also influences CO frequency in females. However, an even lower level of COs is observed in the hybrid (2.79, p = 0.0002), suggesting that an additional phenomenon is responsible for the reduced CO frequency specifically in female hybrids. In all contexts, CO interference is more pronounced in females than males, with the strongest interference observed in female hybrids (Supplemental Figure S9–10). This anti-correlation between CO interference and CO numbers suggests that modulation of CO interference is an important driver of CO numbers. In both sexes of the three backgrounds, CO number is positively correlated with chromosome length, except for the female hybrid where the curve is almost flat at just above 0.5 CO per chromosome per gamete, corresponding to one CO per bivalent and a very strong CO interference (Supplemental Figure S11).

Along chromosomes, a strikingly similar pattern is observed in the three genetic backgrounds. COs are markedly suppressed at the centromeric regions and tend to be frequent at the edge of peri-centromeres in both female and male meiosis. In all three backgrounds, the female and male recombination landscapes tend to diverge with decreasing distance from telomeres, with distal regions being among the highest recombination intervals in males and the lowest in females (Figure 2D–F, Supplemental Figure S12). Strikingly, CO distributions are more closely correlated between the same sexes across the three different backgrounds than between the two sexes in the same background (Figure 2, Supplemental Figure S14). For example, female hybrids are more similar to female Col and L*er* (Spearman’s correlation r_s_ = 0.62 and 0.64) than to male hybrids (r_s_ = 0.26). Thus, sequence divergence appears to have a far lesser impact on the CO landscape than the sex of the meiocyte.

In addition, we wanted to compare the contemporary CO landscapes with the historical landscape. We reconstructed a historical recombination map using a set of non-singleton SNPs generated from 2,029 accessions (Figure 3A and Supplemental Figure S13)^47,48^. Confirming previous findings, the historical CO landscape is strongly correlated with the sequence diversity (Figure 3A). The historical landscape is the result of combined female and male recombination, and we thus compared it to the merged female and male dataset for the inbred lines and to the previously described large Col/L*er* F2 dataset^7^. (Figure 3D–F and Supplemental Figure S14). To facilitate the comparison of the landscapes independently of total CO numbers, we show both the observed CO density (cM/Mb, Figure 3B) and the corresponding normalized distribution (Figure 3A and 3C). Strikingly, the CO landscape in the two inbred lines and hybrid all appear remarkably similar to each other. The distributions of recombination in the inbred and the hybrid genomes closely match each other (Figure 3B), with co-localization of many peaks and valleys, including large peaks on both sides of the centromeric regions, but also in the middle of the arms (Figure 3C). This similarity is confirmed by genome-wide correlation analyses (non-linear correlation r_n_ =0.73–0.78, Figure 3D), which are even stronger when chromosome arms and peri-centromeres are considered separately (Figure 3E-F, r_n_ = 0.78–0.89). The historical recombination landscape is also very similar to the three contemporary landscapes, with most peaks being conserved (r_n_ = 0.6–0.7). This shows that the global CO landscape is largely independent of the presence (hybrid and historical) or absence (Col and L*er*) of polymorphisms between the two chromosomes that recombined. One notable exception is observed at position ∼2 Mb of chromosome 4, with suppression of CO in the hybrid that is not observed in the inbred lines and the historical landscape (Figure 3A–B, 2D–F). This region corresponds to a large ∼1.2 Mb genomic inversion, which suppresses recombination in the Col/L*er* hybrid where it is heterozygous^7,43,49,50^. We then asked if the smaller rearranged regions are also depleted for COs. We explored the overlap of COs with the non-syntenic regions by employing permutation tests in F2 hybrids and observed a strong depletion of CO in non-syntenic regions (Supplemental Figure S15B, p = 0.0002) and CO enrichment in the adjacent regions (Supplemental Figure S15C, p = 0.0002), confirming that structural arrangements are correlated with inhibition of CO formation in hybrids. The CO resolution obtained for inbred lines did not allow us to test if these regions recombine normally in Col and L*er*, as is the case for the unique large inversion.

**Figure 3.**
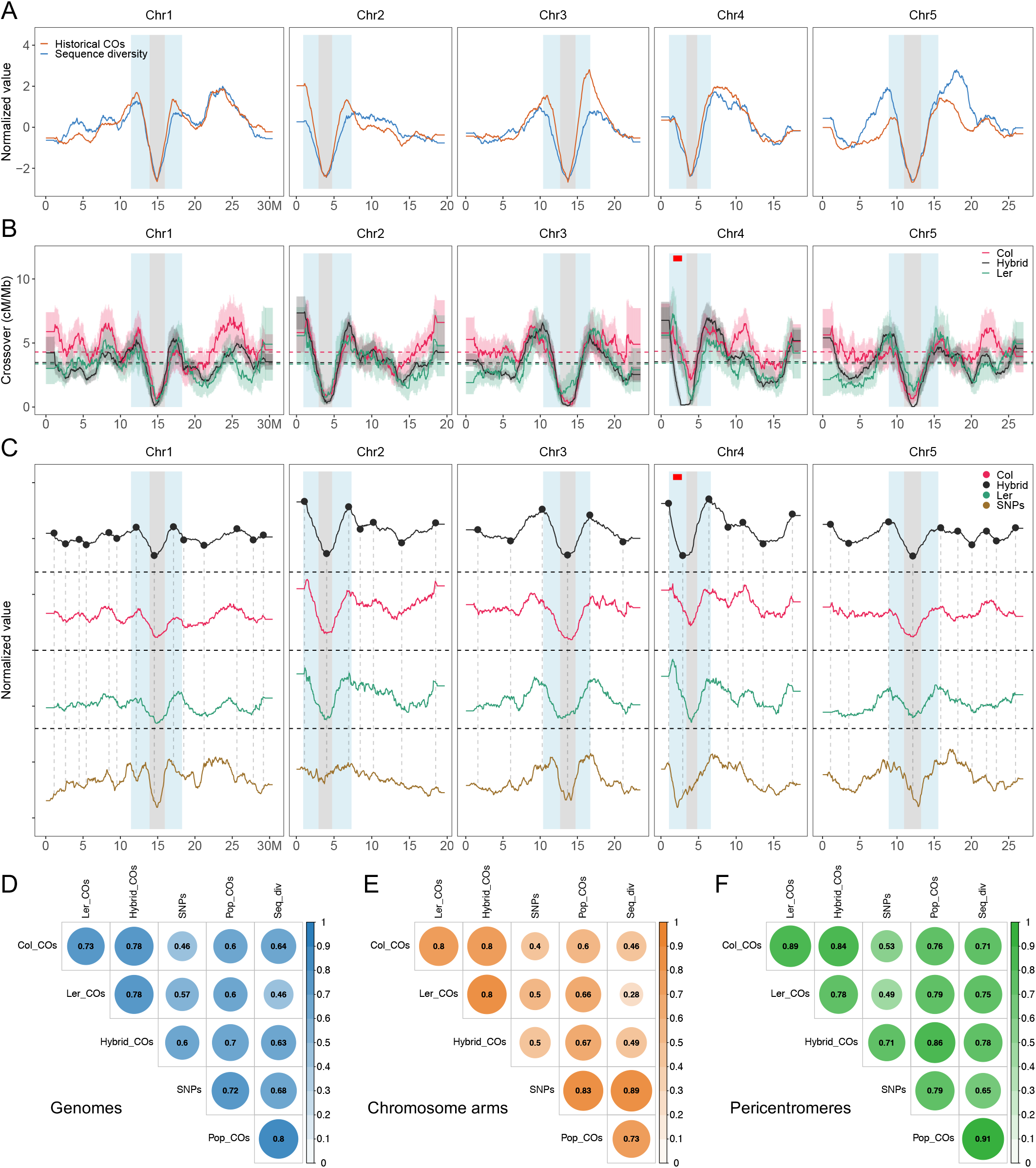
Comparison of the genome-wide CO landscape in Col, L*er*, and F2 hybrids with genetic polymorphisms. (A) The normalized landscapes (sliding window-based, window size 2 Mb and step size 50 kb) of historical recombination rate (4Ner per kb, red) and sequence diversity (π, blue) within 2,029 Arabidopsis accessions across genomes. (B) The chromosomal distribution of COs, with 90% confidence interval in Col, L*er* (merged female and male), and Col/Ler F2 hybrids. The genome-wide mean level recombination is shown with dashed lines. (C) The normalized distribution of COs in Col, L*er*, and F2 hybrids and the density of SNPs between Col and L*er* along chromosomes. The peaks and valleys detected in the hybrid landscape are represented by black points prolonged by vertical dashed lines. The pericentromeric and centromeric regions are indicated by grey and blue shading separately. The ∼1.2 Mb inversion between Col and L*er* on chromosome 4 is represented by a red bar. (D–F) The non-linear correlation coefficient matrices among distributions, considering the (D) whole genome, (E) chromosome arms, and (F) pericentromeric regions. Col_COs, Ler_COs, Hybrid_COs and pop_COs represent CO landscapes in Col, L*er*, F2 hybrids and the population of 2,029 Arabidopsis accessions (historical recombination rate); SNPs represents SNP density between Col and L*er*, and Seq_div represents sequence diversity in the population of 2,029 Arabidopsis accessions.

Consistent with previous analyses^27^, we found that the historical recombination rate is highly correlated with sequence diversity along chromosomes (Figure 3A and 3D–F, Supplemental Figure S14), and contemporary COs in the Col/L*er* hybrid are correlated with SNP density between Col and L*er*^41,51^. As shown above, all the CO landscapes show high similarity. Consequently, the CO landscapes in Col and L*er* are correlated with Col/L*er* SNPs (r_n_ =0.46–0.57) and sequence diversity (r_n_ =0.46–0.64), whereas these polymorphisms were absent in the Col and L*er* inbred lines where these COs were produced. This strongly argues against the possibility that the polymorphisms shape the CO landscape, as the CO landscape is largely unchanged when polymorphisms are absent (with the notable exception of large rearrangements).

To decipher the contributions of genomic and epigenomic features to shaping the CO landscape, we analysed the recombination distribution in Col with a total of 17 different features obtained in the same accession, including GC content; gene and transposable element densities; origins of DNA replication (BrdU-seq)^52^; meiotic DSBs (SPO11-1-oligonucleotides)^53^; chromatin accessibility in flowers (ATAC-seq and DNase-seq)^54–57^; euchromatic (H3K4me1, H3K4me2 and H3K4me3, ChIP-seq)^53,58^ and heterochromatin histone modification marks in flower buds (H3K9me2 and H3K27me1, ChIP-seq)^58^; DNA methylation in male meiocytes (CG, CHG and CHH contexts, BS-seq)^59^; nucleosome occupancy in buds (micrococcal nuclease sequencing, MNase-seq)^53^; and the meiotic cohesin REC8 occupancy (ChIP-seq, Figure 4A, Supplemental Table S2)^58^.

**Figure 4.**
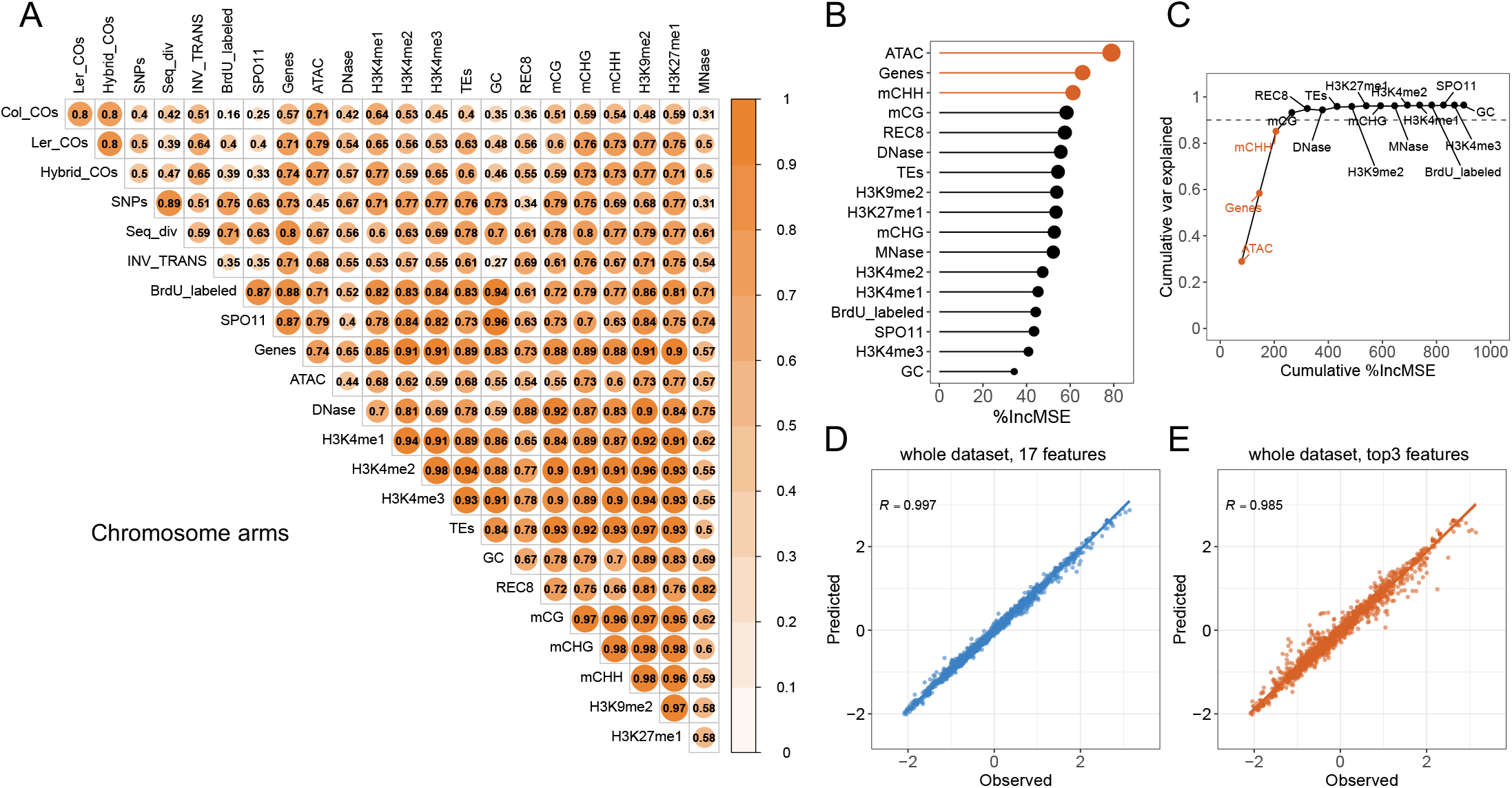
Association and prediction of CO distribution with genomic and epigenomic features. (A) The non-linear correlation matrices show the comparison of pairwise features along chromosome arms, with differences in colour and size according to the correlation scale. Col_COs, Ler_COs and Hybrid_COs (CO landscapes in Col, L*er*, and F2 hybrids), SNPs ( SNPs density between Col and L*er*), Seq_div (sequence diversity in the population of 2,029 Arabidopsis accessions), INV_TRANS (inversions and translocations between Col and L*er*), BrdU_labelled (origins of DNA replication), SPO11 (SPO11-1-oligos), Genes, TEs and GC (gene, TE and GC density), ATAC and DNase (chromatin accessibility, ATAC-seq and Dnase-seq), H3K4me1/2/3, H3K9me2, H3K27me1 (euchromatin, heterochromatin and Polycomb histone marks, ChIP-seq), REC8 (cohesin, ChIP-seq), mCG, mCHG and mCHH (DNA methylation in CG, CHG and CHH contexts), MNase (nucleosome occupancy, MNase-seq). (B) The importance of each of the 17 features for the prediction of CO distribution. The size of points corresponds to the importance. (C) The cumulated proportion of variation that can be explained with the accumulation of features. The top three most important features are colored. (D–E) Spearman’s correlation between the prediction (with the 17 and top three features) and observation of CO distribution, respectively.

Genome-wide, CO distribution is correlated with many genetic and epigenetic features, notably positively with open chromatin (ATAC) (r_n_ =0.71), H3K4me1(r_n_ =0.65), gene density (r_n_ =0.64) and CHG methylation (r_n_ = 0.55) (Supplemental Figure S16A). These correlations are at least partially driven by the centromeric regions, at which COs are abolished and where these features are strongly depleted (Supplemental Figure S16B and S17). However, considering only the chromosome arms, the correlations are almost the same between COs and ATAC (r_n_ = 0.71), H3K4me1(r_n_ = 0.64), CHG methylation (r_n_ = 0.59) gene density (r_n_ = 0.57), and other features (Figure 4A), suggesting a relationship between CO density and chromatin features beyond the centromere.

We next used a machine-learning algorithm (random forest) to analyse the contribution of 17 genomic features on chromosome arms in Col. We first developed a model to predict the frequency of meiotic recombination for a given interval with all the given features together and analysed how the model learned to perform the prediction. ∼96.3% of variation can be explained by the random forest predictive model, as the predicted CO landscape is highly correlated with the observation (Spearman’s r_s_ =0.997, Figure 4D, Supplemental Figure S18A). As shown in Figure 4B and 4C, the most important feature was open chromatin (ATAC), which explained ∼28.7% of the variation. In order to further investigate the performance of the random forest model, we generated a training set by simply sampling 70% of the intervals, with the remaining 30% serving as the testing set. The model trained with the training set performed well with the test set; it could explain ∼94.5% of the variation in the training set (r_s_ =0.996), resulting in a significant correlation (r_s_ =0.977) between the prediction and the observations from the test set (Supplemental Figure S18B, S18E and S18F). Furthermore, we observed that the top three features (open chromatin, gene density, and CHH DNA methylation) can explain ∼85% of the variation (Supplemental Figure S18C). Strikingly, when we trained the model with only these top three features, we observed a high correlation between the predicted and observed CO distribution (r_s_ = 0.985, training set r_s_ = 0.982, and testing set r_s_ = 0.92) (Figure 4E, Supplemental Figure S18D, S18G–H). Altogether, these results show that chromatin accessibility, gene content, and DNA methylation are the major predictors of the mega-scale distribution of meiotic recombination in *Arabidopsis thaliana*.

## Discussion

In this work, we developed a method to analyse the genome-wide recombination landscape in inbred lines and applied it to the Arabidopsis accessions Col and L*er*. This method is based on the introduction of a limited number of markers and allows robust detection of COs. The strategy can be applied to any species for which homozygous lines and mutagenesis are available. We expect this method to be particularly useful for exploring the natural variation of recombination landscapes in species that are inbred lines in the wild (e.g., Arabidopsis) and for exploring CO distribution in species where inter-strain crosses are problematic (e.g., in the fission yeast *Schizosaccharomyces pombe* because of killer meiotic drivers)^60^.

Meiotic recombination frequency has previously been studied in hybrids in many species and varies along chromosomes and positively correlates with the distribution of polymorphisms^3,7,12,21,23–27,51^. One of the possible causes for these correlations is that heterozygosity may favour the formation of COs^21,22^. We showed here that the megabase-scale recombination landscape in inbred lines is strikingly similar to those of hybrids as well as to historical patterns. The observation that the CO landscape is maintained in the absence of polymorphism leads to the conclusion that polymorphisms are not a major determinant of the megabase-scale CO distribution. Polymorphisms, including SNPs and small rearrangements, influence the local recombination pattern^18–21^, but this effect is not manifest at the megabase-scale; at this range, the landscape appears to be unaffected by polymorphism density. An important exception is large-scale genomic rearrangements, such as the ∼1.2 Mb inversion (between Col and L*er*) on chromosome 4, where COs are abolished in hybrids, while the corresponding regions are CO-proficient in isogenic lines. Smaller structural variations are also associated with CO depletion in the hybrid and are presumably CO-prone in the inbred lines, though we cannot confirm this because of the relatively low resolution in CO position.

In many species, COs tend to colocalize with nucleosome- and methylation-depleted gene promoters^3,7,30,34,41,61,62^, consistent with our observation in Col/L*er* hybrids. Moreover, in this study, we found that among a total of 17 genomic and epigenomic features, open chromatin (ATAC), gene density, and DNA methylation in the CHH context are the most potent factors for predicting the distribution of COs along chromosomes in inbred Col. Interestingly, these three features were enough to predict ∼85% of the CO variation along chromosome arms. We do not claim that these three features alone directly control CO positions, but these results show that the chromatin context, which can be largely captured using these three features, efficiently predicts mega-base CO landscapes.

Our results suggest that the large-scale CO landscape is not driven by the polymorphism density. Thus, two possibilities may explain the correlation between polymorphism density and recombination observed in hybrid and historical landscapes. First, the recombination landscape could gradually shape the polymorphism density. Indeed, meiotic recombination is mutagenic, which might be an important driver of genetic diversity and genome evolution^25,28,62–66^. In addition, selection tends to reduce polymorphisms in regions with low recombination rates: both the spread of beneficial mutations and the removal of deleterious mutations by selection reduce polymorphism levels and this effect is larger if recombination is low^67^. A second, not mutually exclusive hypothesis, is that local differences in chromatin features not only influence the distribution of recombination, but that chromatin, independently of recombination, contributes to genomic diversity by shaping differences in local mutation rates along the genome.

While the recombination landscape is largely conserved between the inbred lines and the hybrid, they differ in the total CO number. Globally, there are more COs in Col than in L*er*, with the hybrid having an intermediate number. This is consistent with previous observations in a few crossover reporter intervals and is probably largely driven by an allelic difference in the pro-CO factor HEI10; the Col allele was shown to increase the number of COs compared to the L*er* allele in a co-dominant manner^68^.

When female and male recombination are analysed separately, CO rates are always highest in Col and lowest in L*er* and always higher in males than females. The hybrid exhibits an intermediate recombination rate of CO formation in males, but in contrast the hybrid female has less COs than the two inbred lines. This suggests that some mechanism specifically reduces CO frequency in female hybrids compared to the female inbred lines. One possibility is that class II COs, which represent a minority of COs, are inhibited in the presence of polymorphism and thus reduced in hybrids^49,69^. This would have a proportionally larger effect on female meiosis where class I COs are less numerous than in males, and thus account for the very low level of COs in hybrid females. As class II COs are non-interfering, this would also explain why CO interference is stronger in hybrid females than in inbred females and could especially account for the absence of very close double-COs (Supplemental Figure S10)^70^. Interestingly, in female hybrids, the number of COs observed was very close to the obligate one crossover per bivalent (0.5 crossover per chromatid), suggesting that the CO landscape in female hybrids corresponds to the distribution of the obligate CO, which thus occurs highly preferentially in the proximal regions. The most striking contrast between females and males was the pronounced difference in the distal regions, where males tend to recombine more than females in both pure lines and hybrids. This further confirms that the megabase-scale recombination landscape is largely independent of polymorphisms, and instead suggests that the cellular environment plays a much more critical role, notably by controlling chromosome organization^70,71^.

An improved understanding of the control of meiotic recombination along the chromosome opens the possibility of manipulating COs and increasing recombination rates globally^49,69,72^ and in reluctant regions. This would facilitate the reshuffling of genomic material, breaking of the linkage between beneficial and deleterious alleles, and allow the combination of favourable alleles in elite varieties.

## Materials and methods

### Isogenic population construction and sequencing

Plants were grown in greenhouses or growth chambers (16-h day/8-h night, 20 °C). Wild-type Col-0 and L*er*-1 are 186AV1B4 and 213AV1B1 from the Versailles *Arabidopsis thaliana* stock center (http://publiclines.versailles.inra.fr/). For each accession, seeds were subjected to EMS mutagenesis as described in^73^, and four independent M2 plants were crossed to produce two independent F1*, which were consequently heterozygous for a set of EMS mutations (Figure 1). Then, the two F1* plants were reciprocally crossed to generate two F1 populations. To test the robustness of the results and detect the unlikely possibility that a dominant modifier of recombination was caused by an EMS-induced mutation, two independent replicates of the entire process were performed for each accession. These F1* and F1 plants were then used for CO analysis by whole-genome sequencing (Figure 1, Supplemental Table S1). Leaf samples from the populations were used for DNA purification and library preparation for 2×150bp HiSeq 3000 Illumina sequencing^74^. To detect the markers, we sequenced genomic DNA from the F1* (∼59 × and ∼16x, in Col and L*er* respectively) and F1s (∼4.8x and ∼5.0x)

### Identifying and genotyping EMS-induced mutations

For each individually sequenced F1* and F1 plant of *Arabidopsis* Col and L*er* accessions, the whole-genome resequencing reads were aligned against the Col-0 TAIR10 reference genome^75,76^ and L*er* assembled genome^43^ by BWA v0.7.15-r1140^77^ with default parameters, and variant calling in F1* populations was performed using inGAP-family^37^, separately. To obtain high-quality mutation marker lists, we first removed non-allelic markers using inGAP-family with input from the tandem replicates and structural variants predicted using Tandem Repeats Finder v4.09^78^ and inGAP-family, respectively, and further filtered variations that not meet the following criteria: (i) heterozygous genotype with alternative allele frequency from 0.4 to 0.6, (ii) specific to each of the F1*, and (iii) GA to CT substitution. Then, the read count and genotype map of mutation markers of each F1* was generated from their F1 progenies by inGAP-family, which was subsequently used for mutation phasing and CO identification. In order to properly compare CO landscapes in isogenic and hybrid lines, we transferred the coordinates of mutations in the L*er* population to Col-0 by using syntenic alignments identified by SyRI v1.2^79^.

Two additional replicates in Col (C and D) were discarded, because the marker analysis showed that one of the F1* was resulting from an accidental selfing and not from a cross. Two additional replicates in L*er* were also discarded (C and H), because the number of detected mutation markers (<350) was insufficient for good genome coverage.

### Phasing mutations and CO identification in inbred lines

To phase the EMS-induced mutation markers, we employed a hierarchical clustering-based sliding window method, with a window size of ten mutation markers and step size of one mutation marker (Supplemental Figure S1). For each window, the genotype map of the mutation markers was constructed and used as input for clustering, resulting in two groups: one consisting of wild-type samples and one comprising mutant samples. The genotype and phase of mutation was evaluated by the vote of covered sliding windows. During this process, uncovered and low-covered markers were imputed and corrected. The CO events were defined as consistent switches of phase of mutation markers along chromosome arms, and the border was further refined by examining the wild-type allele of the mutation. For the termini of chromosomes, COs were validated as switches with one well-supported mutant allele or more than ten reads supported by the consecutive wild-type allele of the variant marker. The CO interference was analyzed using MADpattern^80,81^.

### CO analysis in hybrid population

The Col/L*er* and wild-type Col plants were reciprocally crossed to construct female and male populations (428 and 294 plants, respectively). Leaf samples of the backcross populations were collected for DNA purification, library preparation, and Illumina sequencing^74^. In addition, the raw reads of the Col/L*er* F2 population were downloaded from ArrayExpress with the accession number E-MTAB-8165^7^.

The quality of the raw sequencing datasets was checked using FastQC v0.11.9 (http://www.bioinformatics.babraham.ac.uk/projects/fastqc/), and then adapters and low-quality bases were trimmed using Trimmomatic v0.38^82^, with parameters “LEADING:5 TRAILING:5 SLIDINGWINDOW:5:20 MINLEN:50”. The mapping of sequencing reads, generation of high-confidence SNP markers between Col and L*er*, meiotic CO prediction, and filtering of the poorly covered and potentially contaminated samples were performed using a previously described method^44^. To avoid false genotyping, we selected 0.95 as the threshold allelic ratio for the determination of homozygosity in F1 hybrids. Identified COs were manually checked at random using inGAP-family.

### SPO11-1-oligo, BrdU-seq, ChIP-seq, MNase-seq, DNase-seq, and ATAC-seq data analysis

The short reads from public datasets (Supplemental Table S2) were quality-checked with FastQC. The specific 3’ adapter and 5’ ends sequences were trimmed before alignment by Cutadapt v1.9.1^83^ as described^53^. For BrdU-seq and ATAC-seq datasets, the reads were processed with Trimmomatic to remove potential adapter sequences and low-quality bases, with “LEADING:3 TRAILING:3 SLIDINGWINDOW:4:15 MINLEN:36”. Duplicated reads were removed using BBMap (https://github.com/BioInfoTools/BBMap). Then, clean reads were aligned to the TAIR10 reference genome using Bowtie2 v2.2.8^84^ with settings “--very-sensitive -k 10” for single-end datasets and further settings “--no-discordant --no-mixed” for paired-end datasets. The uniquely mapped reads were kept for subsequent analysis, which were processed by Samtools v1.9^85^ and Sambamba v0.6.8^86^. For all sequencing data, coverage across the genome was evaluated and normalized with bins per million mapped reads (BPM) in bedGraph format using bamCoverage v3.4.3^87^.

### Bisulfite sequencing data analysis

The quality and adapter sequencing of raw reads were examined by FastQC. The sequencing reads were mapped to the TAIR10 reference genome with Bismark v0.22.0^88^, with the following setting: -q -bowtie2 -N 1 -L 24. Reads that mapped to multiple positions and duplicated alignments were removed. Methylated cytosines in the CG, CHG, and CHH contexts, and the level of methylation were extracted for subsequent association analysis.

### Genome-wide CO distribution correlation analysis

The chromosomal profiles of COs, genomic, and epigenomic features were estimated in 50kb windows along chromosomes. For a given window, the recombination frequency was normalized with the total CO number within the corresponding chromosome. Then, all of the COs, genomic and epigenomic data were smoothed with 40 nearby windows using the filter function and then normalized using the scale function in the R environment. The non-linear correlation matrices were calculated using the nlcor package (https://github.com/ProcessMiner/nlcor) in R, at the genome, chromosome arms, and pericentromeric scale respectively. The constitution (pericentromeres, centromeres, and arms) of the TAIR10 reference genome was adopted from Underwood et al.^61^. Here, all the random forest models were trained using randomForest v4.6-14 package in R, with the setting of “mtry=3, importance=TRUE, na.action=na.omit, ntree=2000”.

### Estimating nucleotide diversity and historical recombination rate

For the sequence polymorphism data of 2,029 Arabidopsis accessions from the 1,001 Genomes Project^47^ and the RegMap population^48^, we first selected diallelic SNP positions with <20% missing data and >5% minimum allele frequency using VCFtools v0.1.16^89^. Then, we masked SNPs located in (i) tandem repeat regions (Tandem Repeats Finder output), (ii) repetitive elements and low-complexity regions (extracted from the masked TAIR10 reference genome), (iii) transposable elements (TAIR10 annotation) and (iv) centromeric regions (definition adopted from Underwood et al.^61^. Finally, we obtained a collection of 905,613 SNPs from 2,029 accessions for CO frequency analysis. FastEPRR v2.0^90^ was employed for estimating population recombination rates (ρ = 4Ner, where Ne is the effective population size and r is the recombination rate of the window), with 50 kb non-overlapping window size. The nucleotide diversity of each 50 kb non-overlapping window along chromosomes was calculated using VCFtools.

## Data availability

The raw sequencing data of individuals of Col, L*er* inbred lines and the Col/L*er* F1 hybrid can be accessed in ArrayExpress under the accession numbers E-MTAB-11248, E-MTAB-11249, E-MTAB-11250, E-MTAB-11251, E-MTAB-11254, respectively. The list of COs identified in Col, L*er*, F1 hybrid (female and male), and F2 hybrid can be found in Supplemental Table S3-S6.

## Acknowledgements

We would like to thank the Max Planck Genome center for DNA extraction, library preparation and sequencing, Hequan Sun and Wen-Biao Jiao for helpful discussion, Charles Underwood, Ian Henderson and Andrew Tock for the help with SPO11-oligos analysis. We thank Wayne Crismani and Andrew Lloyd for critical reading of the manuscript.

This work was support by core funding from the Max Planck Society, and an Alexander von Humboldt Fellowship to Q.L.

## Author contributions

Q.L., K.S. and R.M. designed the research. Q.L., K.S. and R.M. analyzed the data. V.S. and B.W. generated plant materials. B.H. supervised the whole-genome sequencing work. Q.L. and R.M. wrote the article with input from K.S.

## Supplemental Figure legends

**Figure S1. Mutation marker phasing and CO identification.**

The EMS-induced mutation markers were first genotyped in each F1 population. Then, in each sliding window of markers, F1 samples were clustered into two groups, corresponding to the two different phases (P1 and P2, alleles coloured by blue and orange). After phasing, the count table was generated, indicating the number of supported windows of each phase on each marker position. With this, uncovered and mis-genotyped markers were imputed and corrected. Finally, COs can be identified as consistent switches of phase. “G” and “A” represent markers that genotyped as reference and mutant alleles, “n” represent markers not covered by sequencing reads.

**Figure S2. The chromosomal distribution of phased EMS-induced mutation markers in Col and L*er*.**

**Figure S3. The phased genotype map of mutation markers on chromosome 1 in Col A population.**

Each line is a F1 individual, each column a marker. As the F1 results from reciprocal crosses between independent F1*, a marker in a given F1* can be either heterozygous or wild-type. Dark purple and dark red indicate the detection of mutant reads (phase 1 and phase 2, respectively), unambiguously scored as a heterozygote. When only wild-type reads are detected, the number of reads is indicated with a colour scale. (phase 1 and phase 2 are colored blue and orange, respectively. Maximum was set as 5 manually). The identified COs are marked with pairs of black arrows.

**Figure S4. The genome-wide phase map of mutation markers in two samples of Col A population.**

**Figure S5. Correlation analysis of CO numbers, sequencing depths and number of covered and phased mutation markers in each replicate population of Col and L*er*.**

**Figure S6. The distribution of interval length of COs and correlation analysis with sequencing depths in Col, L*er*, and hybrids.**

**Figure S7. The chromosomal distribution of COs in Col/L*er* F1 hybrids and relationship to genome features.**

(A) Distribution of COs in female meiosis of Col/L*er* F1 hybrids in this study (pink) compared with Giraut et al (black). (B) Distribution of COs in male meiosis of Col/L*er* F1 hybrids in this study (blue) compared with Giraut et al (black). The exactly same wind size is adopted from Giraut et al for comparison. (C) Sliding window-based distribution (window size 500 kb, step size 50 kb) of COs in female and male meiosis of Col/L*er* F1 hybrids (this study). (D) Distribution of distance of COs from nearest TSS. (E) Permutation tests (5,000 shuffling) were performed to evaluate the overlap between promoter regions of proteins encoding genes and COs in Col/L*er* F1 hybrids.

**Figure S8. The distribution of CO frequency across genomes in Col/Ler detected in backcrosses (F1, this study) and in selfing (F2, Rowan et al) populations.** Comparison was done using a window size of 500 kb and step size of 50 kb (A) or a window size of 2 Mb and step size of 50 kb (B). Female and male data (this study, blue) were merged to be comparable with the selfing data (Rowan et al., orange).

**Figure S9. Comparison of the CoC curve between female and male meiosis in Col, L*er*, and F1 hybrids.**

Chromosomes were divided into 15 intervals, and the mean coefficient of coincidence (CoC = observed number of coincident occurrence of COs in both intervals/expected number according to CO frequency in each interval) was calculated for pairs of intervals separated by a certain distance (proportion of chromosome length). A CoC lower than 1 indicates the presence of crossover interference.

**Figure S10. Comparison of the CoC curve between Col, L*er*, and F1 hybrids in female and male meiosis.**

Chromosomes were divided into 15 intervals, and the mean coefficient of coincidence (CoC = observed number of coincident occurrence of COs in both intervals/expected number according to CO frequency in each interval) was calculated for pairs of intervals separated by a certain distance (proportion of chromosome length). A CoC lower than 1 indicates the presence of crossover interference.

**Figure S11. CO number distribution and correlation analysis with chromosome length in female and male Col, L*er*, and hybrid individuals.**

(A) Distribution of CO numbers per chromosome and per gamete in female and male meiosis of Col, L*er*, and F1 hybrids. (B) Mean number of COs per chromosome per gamete versus chromosome size in Mb, in female and male Col, L*er*, and F1 hybrids. Pearson’s correlation analysis of relationship between CO number and chromosome length: Col female r = 0.96, P = 0.008, Col male = 0.96, P = 0.011, L*er* female r = 0.97, P = 0.007, L*er* male r = 0.94, P = 0.017, Hybrid female r = 0.71, P = 0.18, Hybrid male r = 0.97, P = 0.007. (C) Pearson’s correlation analysis of relationship between CO number and chromosome length in Col, L*er* pseudo F2, and hybrids F2 (Rowan et al) (Col r = 0.96, P = 0.01, L*er* r = 0.98, P = 0.003, Hybrid r = 0.98, P = 0.004).

**Figure S12. Comparison of the chromosomal distribution of COs between Col, L*er*, and F1 hybrids in female and male meiosis.**

The sliding window-based distribution (window size 2 Mb, step size 50 kb) of COs in female (A) and male (B) meiosis of Col, L*er*, and hybrids individually.

**Figure S13. Historical recombination rate in the *Arabidopsis thaliana* genome.**

(A) The geographical distribution of the 2,029 Arabidopsis accessions (the 1001 Genome Project and the RegMap lines, the latitude and longitude for eight accessions were not included). (B) The distribution of historical recombination rates (4Ner per kb) along chromosomes at 50kb window scale within 2,029 Arabidopsis accessions. Mean values are indicated by horizontal lines. The pericentromeric and centromere regions are indicated by grey and blue shading separately.

**Figure S14. Correlation analysis of CO distribution with polymorphisms.**

Spearman’s correlation between each pair of features is shown in the upper triangle panel.

**Figure S15. Permutation analysis of the overlap between non-syntenic and adjacent syntenic regions and COs in Col/L*er* F2 hybrids.**

(A) The percentages of non-syntenic and syntenic regions in Col and L*er* and pericentromeric regions. (B-C) Permutation tests were performed to evaluate the overlap between non-syntenic and adjacent syntenic regions and COs in Col/L*er* F2 hybrids, respectively. The adjacent syntenic regions were defined as the up- and downstream regions (same size) of non-syntenic regions. The vertical black dashed lines indicate the expected mean number of overlaps in 5,000 shuffled permutations. The observed number of overlaps is represented by vertical red dashed lines.

**Figure S16. Non-linear correlation coefficient matrices for the distribution of COs, genomic and epigenomic features at genome and pericentromeric scales.**

Col_COs, Ler_COs and Hybrid_COs (CO landscapes in Col, L*er*, and F2 hybrids), SNPs ( SNPs density between Col and L*er*), Seq_div (sequence diversity in the population of 2,029 Arabidopsis accessions), INV_TRANS (inversions and translocations between Col and L*er*), BrdU_labelled (origins of DNA replication), SPO11 (SPO11-1-oligos), Genes, TEs and GC (gene, TE and GC content density), ATAC and DNase (chromatin accessibility, ATAC-seq and Dnase-seq), H3K4me1/2/3, H3K9me2, H3K27me1 (euchromatin, heterochromatin and Polycomb histone marks, ChIP-seq), REC8 (cohesin, ChIP-seq), mCG, mCHG and mCHH (DNA methylation in CG, CHG and CHH contexts), MNase (nucleosome occupancy, MNase-seq).

**Figure S17. Genomic landscape of Col COs, genomic and epigenomic features.**

Sliding window-based distributions (window size 2 Mb, step size 50 kb) were normalized to the same scale. The pericentromeric and centromere regions are indicated by grey and blue shading separately. Col (CO landscapes in Col), BrdU (origins of DNA replication), SPO11 (SPO11-1-oligos), Genes and TEs (gene and TE density), ATAC and DNase (chromatin accessibility, ATAC-seq and Dnase-seq), H3K4me1 and H3K9me2 (euchromatin and heterochromatin histone marks, ChIP-seq), REC8 (cohesin, ChIP-seq), mCG, mCHG and mCHH (DNA methylation in CG, CHG and CHH contexts), MNase (nucleosome occupancy, MNase-seq).

**Figure S18. The performance of the random forest model for predicting CO distribution in Col.**

(A-B) The distribution of observed and predicted CO frequency by modelling with 17 features in the whole, training (70%) and testing (30%) datasets, respectively. (C-D) The distribution of observed and predicted CO frequency by modelling with the top three most important features in the whole, training and testing datasets, respectively. The pericentromeric and centromere regions are indicated by grey and blue shading separately. (E-H) Spearman’s correlation between the predicted and observed CO distribution predicted by models constructed with 17 features and the top three most important features in the training and testing datasets, respectively.

**Table S1. Summary of EMS-induced mutations in Arabidopsis inbred lines Table S2. The public dataset used in this study**

**Table S3. CO positions in inbred Col**

**Table S4. CO positions in inbred L*er***

**Table S5. CO positions in Col × L*er* backcross population**

**Table S6. CO positions in Col × L*er* F2 population**

