## supplemental figures and tables for "The megabase-scale crossover landscape is independent of sequence divergence"

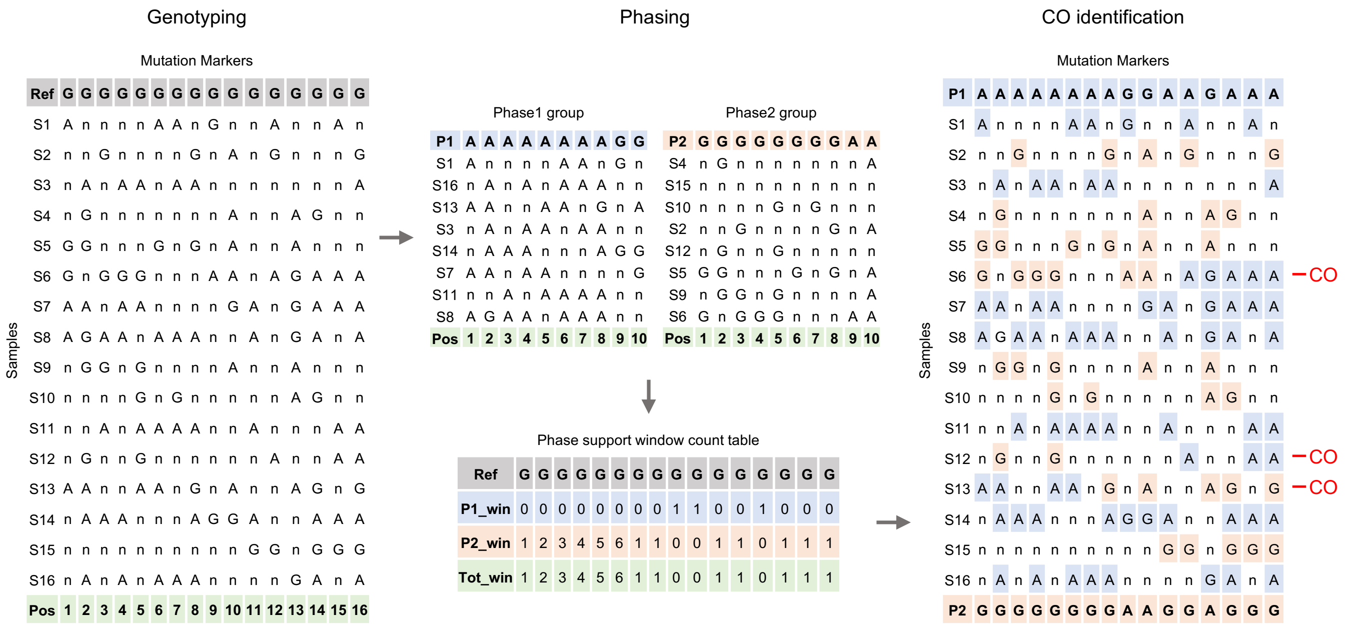


**Figure S1. Mutation marker phasing and CO identification.**
The EMS-induced mutation markers were first genotyped in each F1 population. Then, in each sliding window of markers, F1 samples were clustered into two groups, corresponding to the two different phases (P1 and P2, alleles coloured by blue and orange). After phasing, the count table was generated, indicating the number of supported windows of each phase on each marker position. With this, uncovered and mis-genotyped markers were imputed and corrected. Finally, COs can be identified as consistent switches of phase. “G” and “A” represent markers that genotyped as reference and mutant alleles, “n” represent markers not covered by sequencing reads.

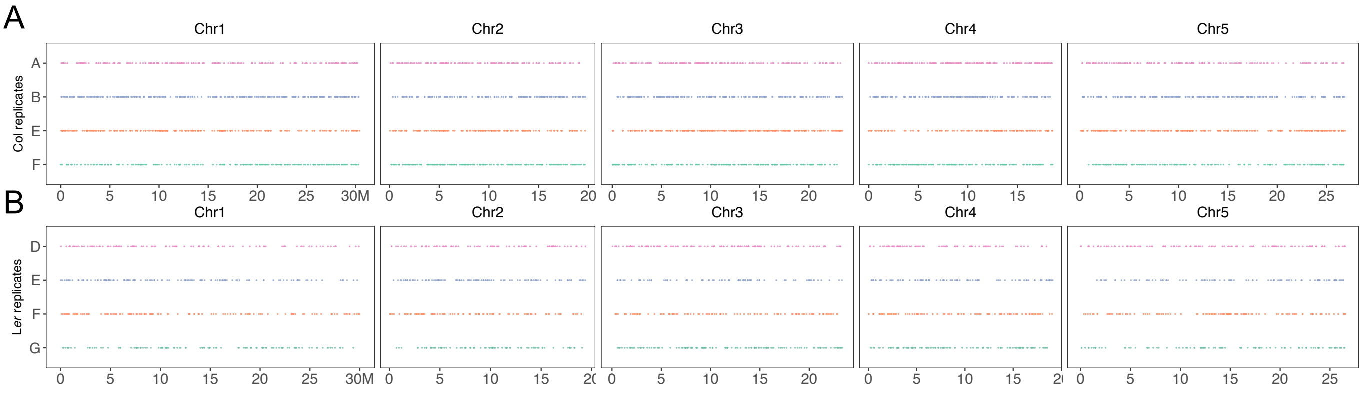


**Figure S2. The chromosomal distribution of phased EMS-induced mutation markers in Col and L*er*.**

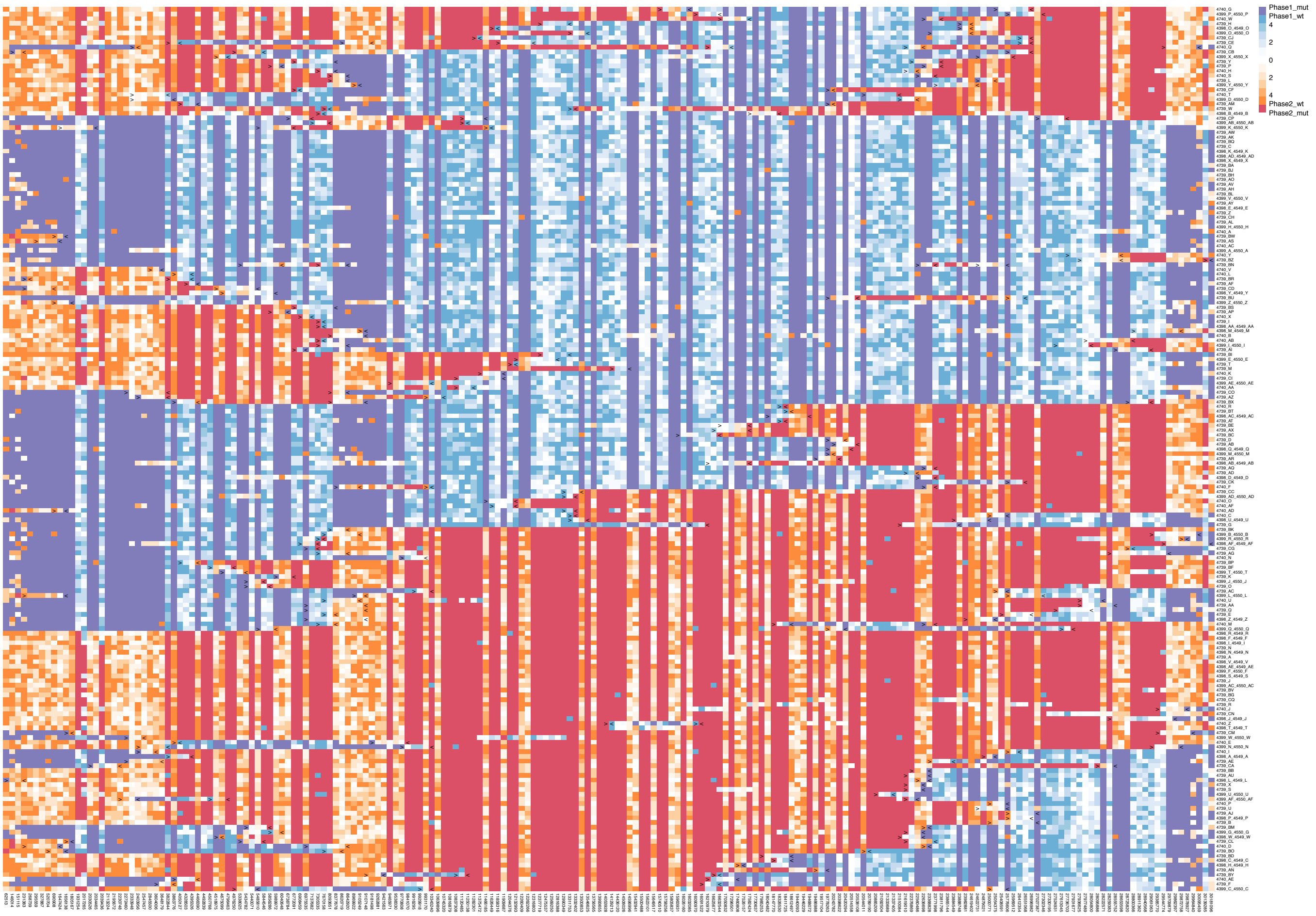


**Figure S3. The phased genotype map of mutation markers on chromosome 1 in Col A population.** Each line is a F1 individual, each column a marker. As the F1 results from reciprocal crosses between independent F1*, a marker in a given F1* can be either heterozygous or wild-type. Dark purple and dark red indicate the detection of mutant reads (phase 1 and phase 2, respectively), unambiguously scored as a heterozygote. When only wild-type reads are detected, the number of reads is indicated with a colour scale. (phase 1 and phase 2 are colored blue and orange, respectively. Maximum was set as 5 manually). The identified COs are marked with pairs of black arrows.

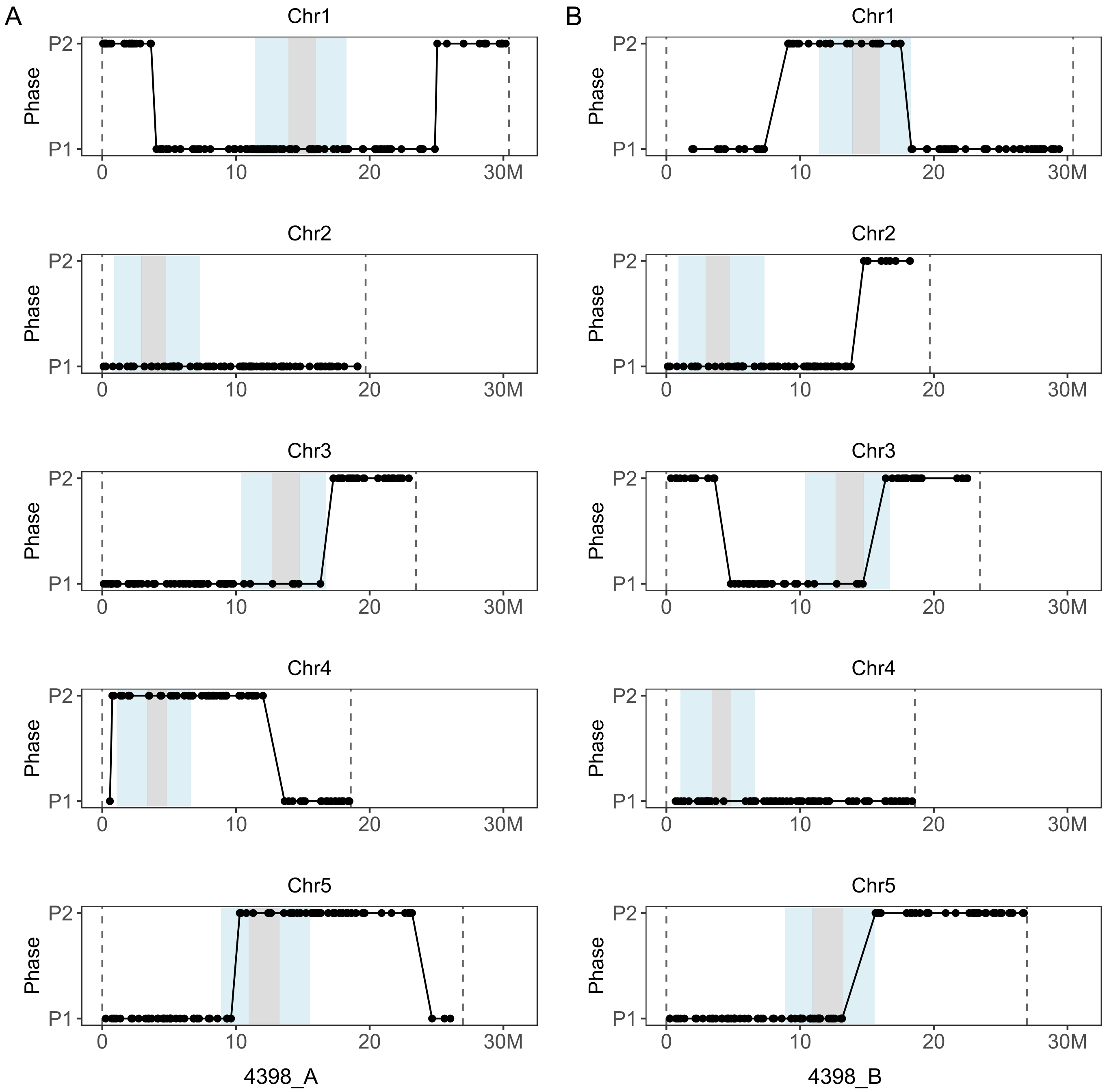


**Figure S4. The genome-wide phase map of mutation markers in two samples of Col A population.**

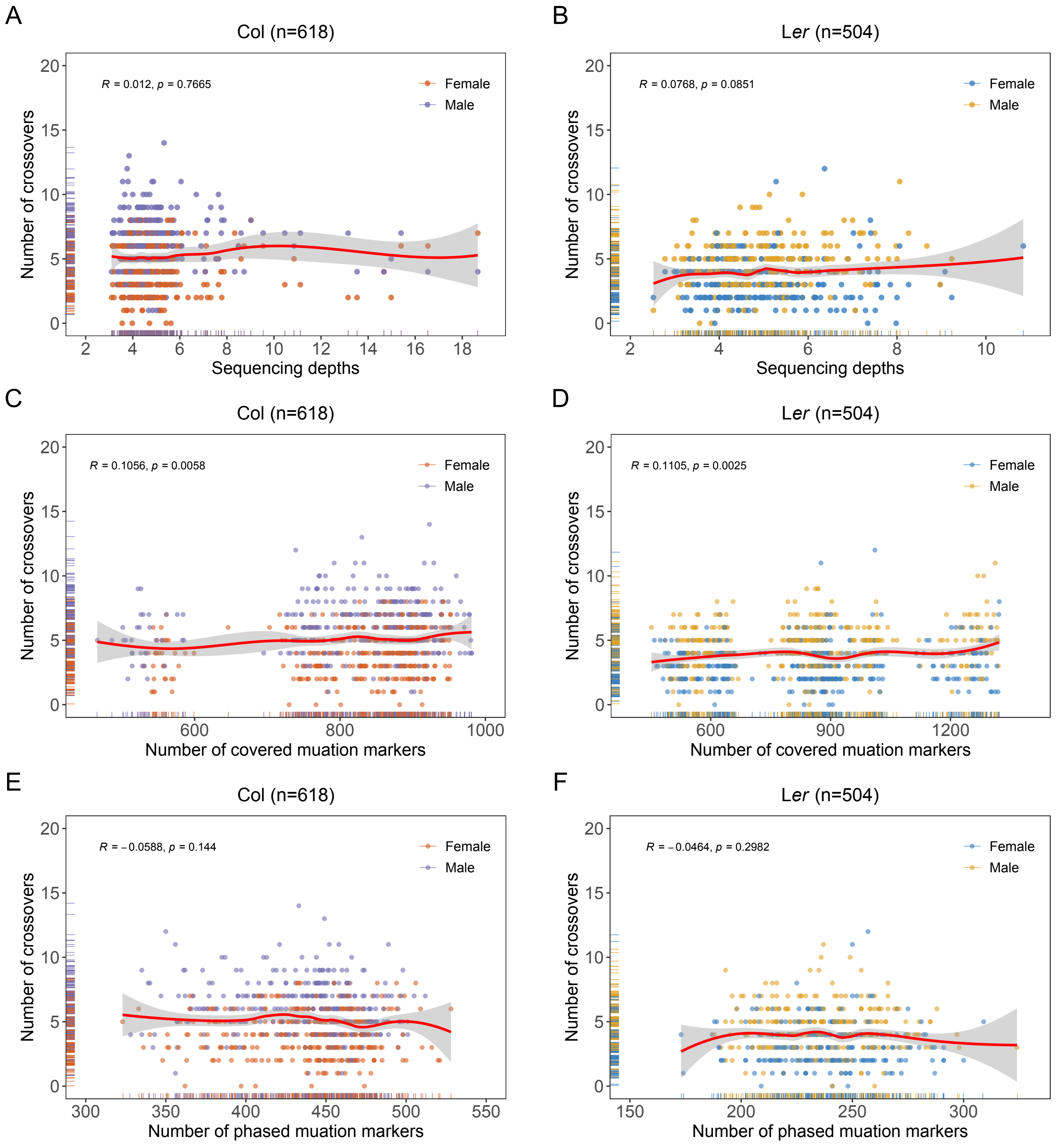


**Figure S5. Correlation analysis of CO numbers, sequencing depths and number of covered and phased mutation markers in each replicate population of Col and L*er*.**

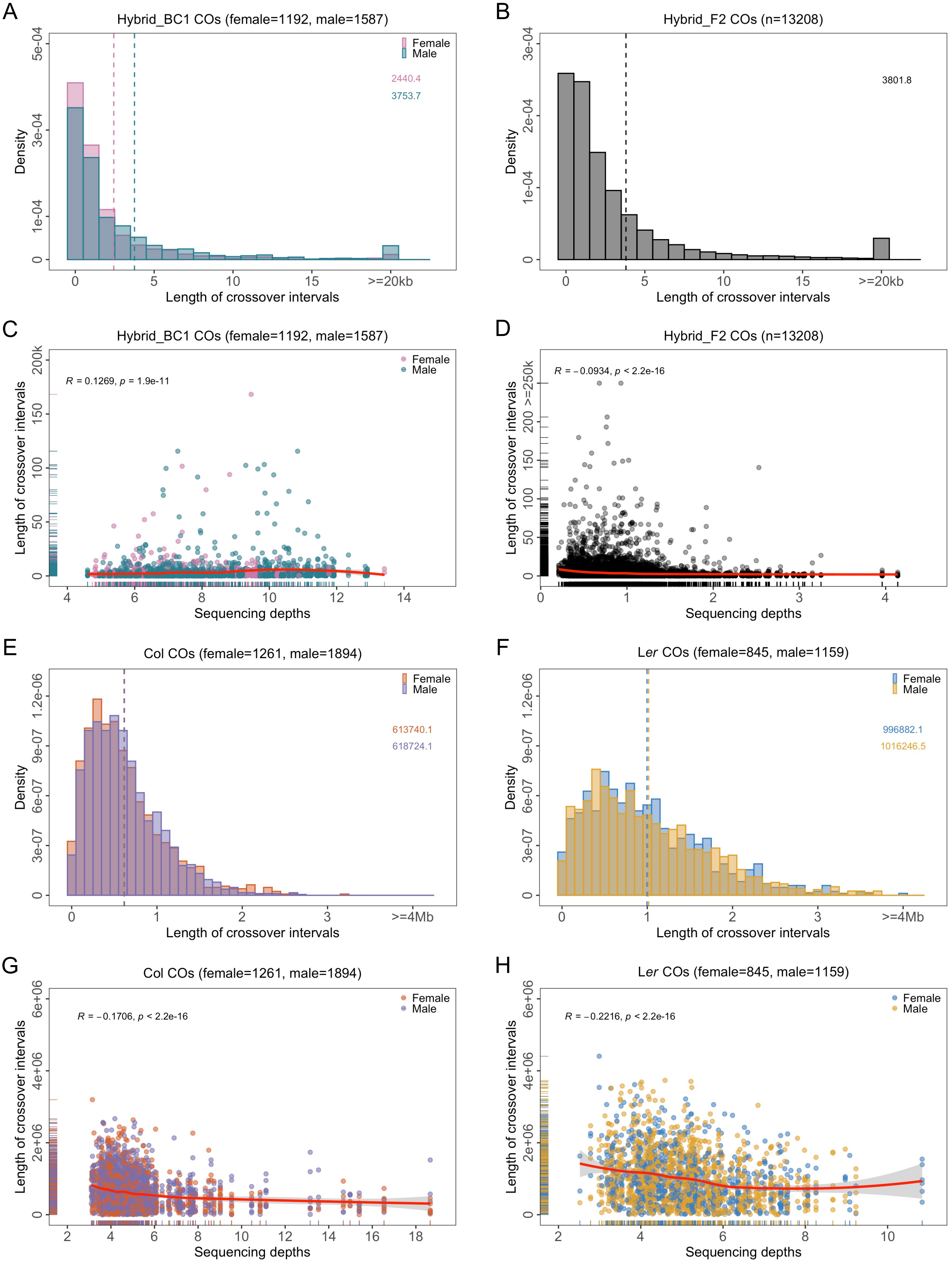


**Figure S6. The distribution of interval length of COs and correlation analysis with sequencing depths in Col, L*er*, and hybrids.**

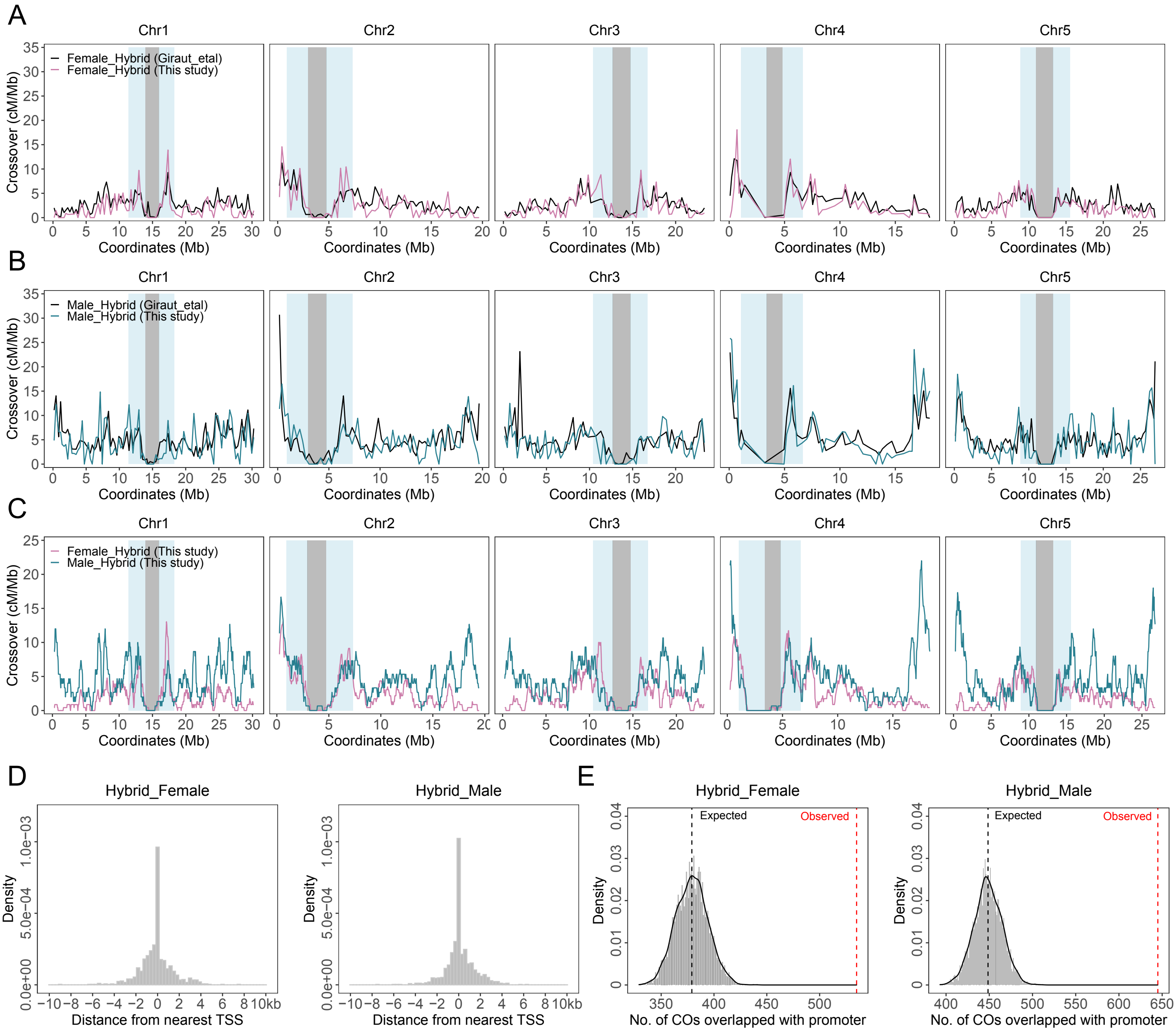


**Figure S7. The chromosomal distribution of COs in Col/L*er* F1 hybrids and relationship to genome features.**

(A) Distribution of COs in female meiosis of Col/L*er* F1 hybrids in this study (pink) compared with Giraut et al (black). (B) Distribution of COs in male meiosis of Col/L*er* F1 hybrids in this study (blue) compared with Giraut et al (black). The exactly same wind size is adopted from Giraut et al for comparison. (C) Sliding window-based distribution (window size 500 kb, step size 50 kb) of COs in female and male meiosis of Col/L*er* F1 hybrids (this study). (D) Distribution of distance of COs from nearest TSS. (E) Permutation tests (5,000 shuffling) were performed to evaluate the overlap between promoter regions of proteins encoding genes and COs in Col/L*er* F1 hybrids.

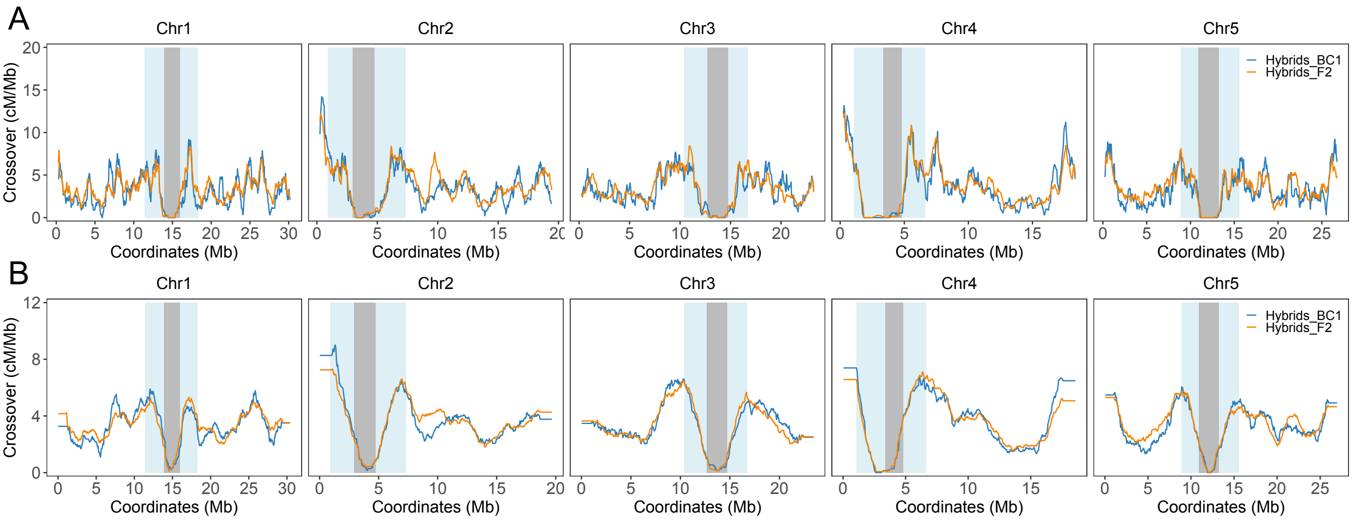


**Figure S8. The distribution of CO frequency across genomes in Col/L*er* detected in backcrosses (F1, this study) and in selfing (F2, Rowan et al) populations.**

Comparison was done using a window size of 500 kb and step size of 50 kb (A) or a window size of 2 Mb and step size of 50 kb (B). Female and male data (this study, blue) were merged to be comparable with the selfing data (Rowan et al., orange).

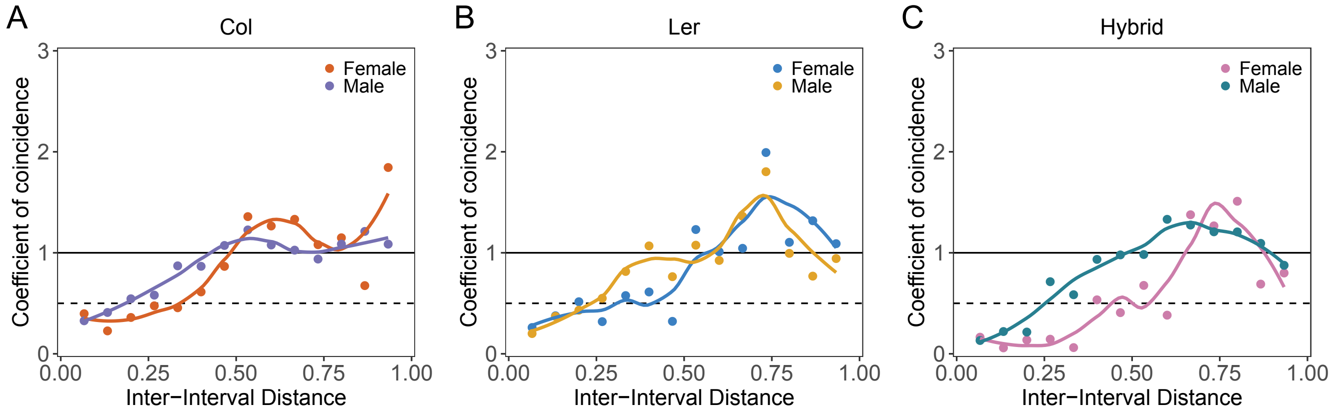


**Figure S9. Comparison of the CoC curve between female and male meiosis in Col, L*er*, and F1 hybrids.** Chromosomes were divided into 15 intervals, and the mean coefficient of coincidence (CoC = observed number of coincident occurrence of COs in both intervals/expected number according to CO frequency in each interval) was calculated for pairs of intervals separated by a certain distance (proportion of chromosome length). A CoC lower than 1 indicates the presence of crossover interference.

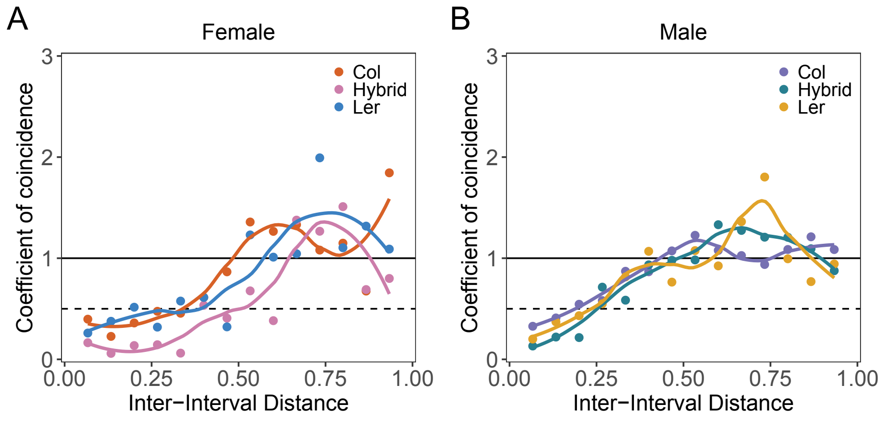


**Figure S10. Comparison of the CoC curve between Col, L*er*, and F1 hybrids in female and male meiosis.** Chromosomes were divided into 15 intervals, and the mean coefficient of coincidence (CoC = observed number of coincident occurrence of COs in both intervals/expected number according to CO frequency in each interval) was calculated for pairs of intervals separated by a certain distance (proportion of chromosome length). A CoC lower than 1 indicates the presence of crossover interference.

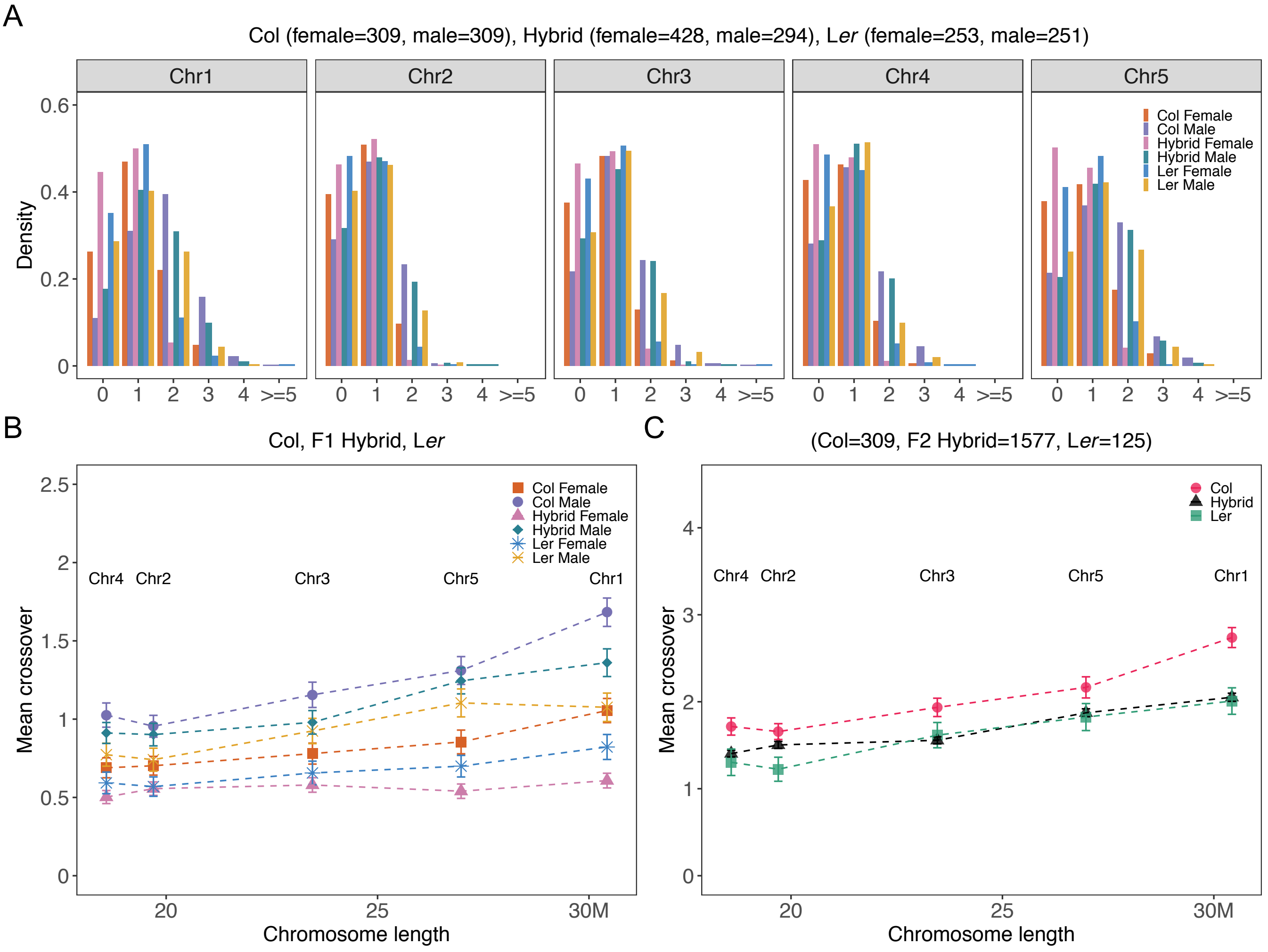


**Figure S11. CO number distribution and correlation analysis with chromosome lengths in female and male of Col, L*er*, and hybrids.**

(A) Distribution of CO numbers per chromosome and per gamete in female and male meiosis of Col, L*er*, and F1 hybrids. (B) Mean number of COs per chromosome per gamete versus chromosome size in Mb, in female and male Col, L*er*, and F1 hybrids. Pearson's correlation analysis of relationship between CO number and chromosome length: Col female r = 0.96, P = 0.008, Col male = 0.96, P = 0.011, L*er* female r = 0.97, P = 0.007, L*er* male r = 0.94, P = 0.017, Hybrid female r = 0.71, P = 0.18, Hybrid male r = 0.97, P = 0.007. (C) Pearson's correlation analysis of relationship between CO number and chromosome length in Col, L*er* pseudo F2, and hybrids F2 (Rowan et al) (Col r = 0.96, P = 0.01, L*er* r = 0.98, P = 0.003, Hybrid r = 0.98, P = 0.004).

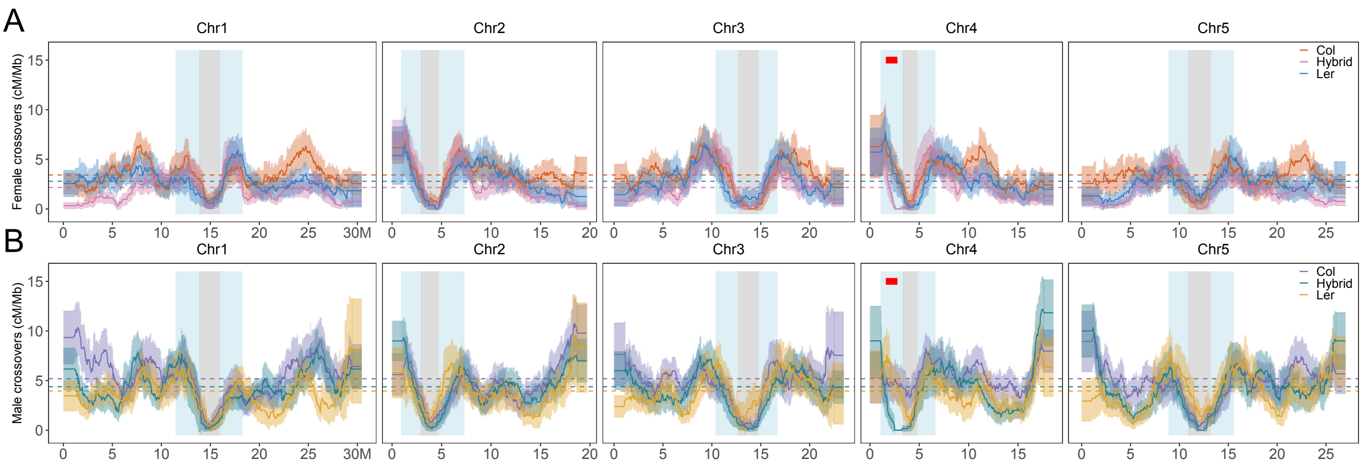


**Figure S12. Comparison of the chromosomal distribution of COs between Col, L*er*, and F1 hybrids in female and male meiosis.**

The sliding window-based distribution (window size 2 mb, step size 50 kb) of COs in female (A) and male (B) meiosis of Col, L*er*, and hybrids individually.

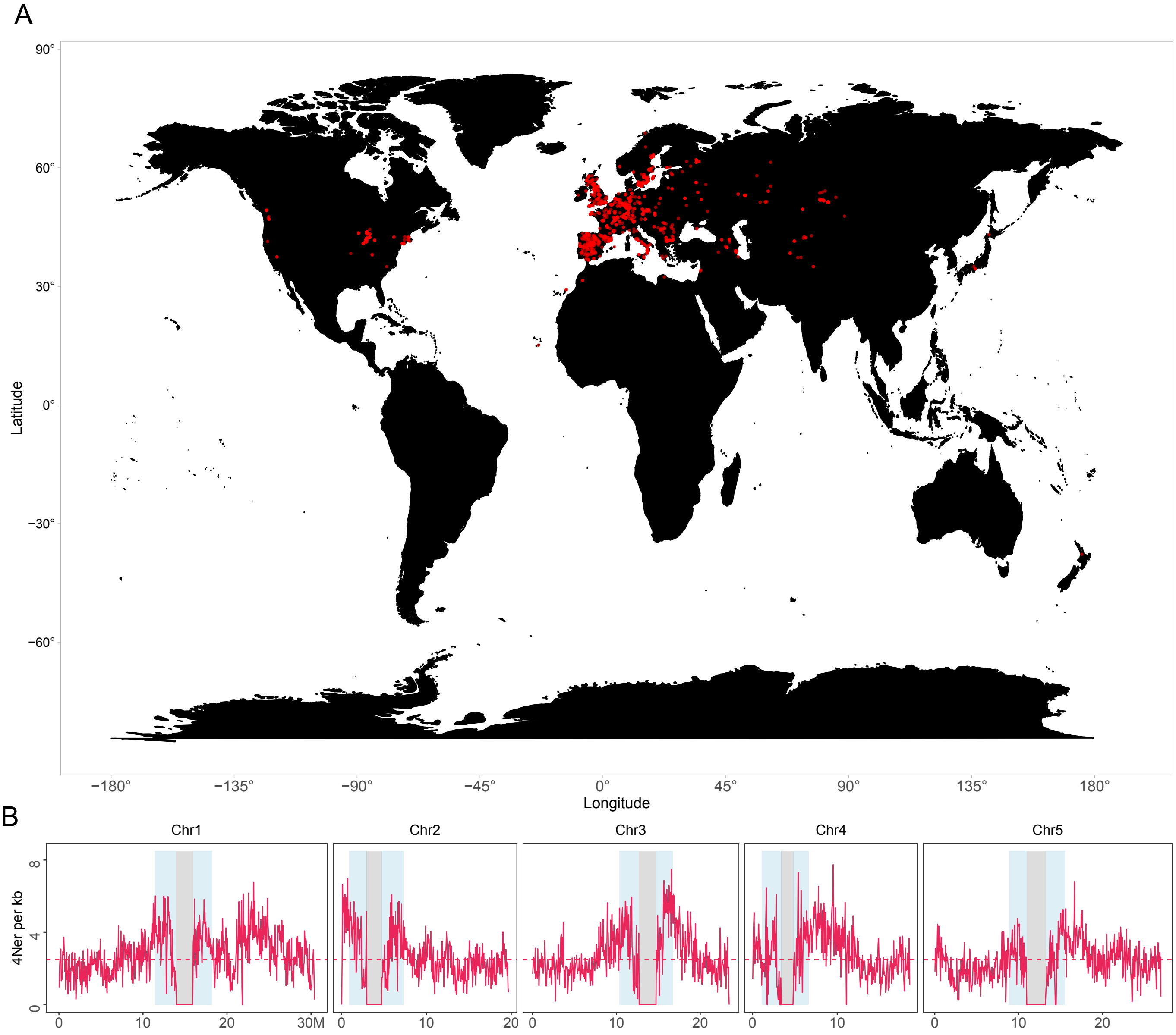


**Figure S13. Historical recombination rate in the *Arabidopsis thaliana* genome.**
(A) The geographical distribution of the 2,029 Arabidopsis accessions (the 1001 Genome Project and the RegMap lines, the latitude and longitude for eight accessions were not included). (B) The distribution of historical recombination rates (4Ner per kb) along chromosomes at 50kb window scale within 2,029 Arabidopsis accessions. Mean values are indicated by horizontal lines. The pericentromeric and centromere regions are indicated by grey and blue shading separately.

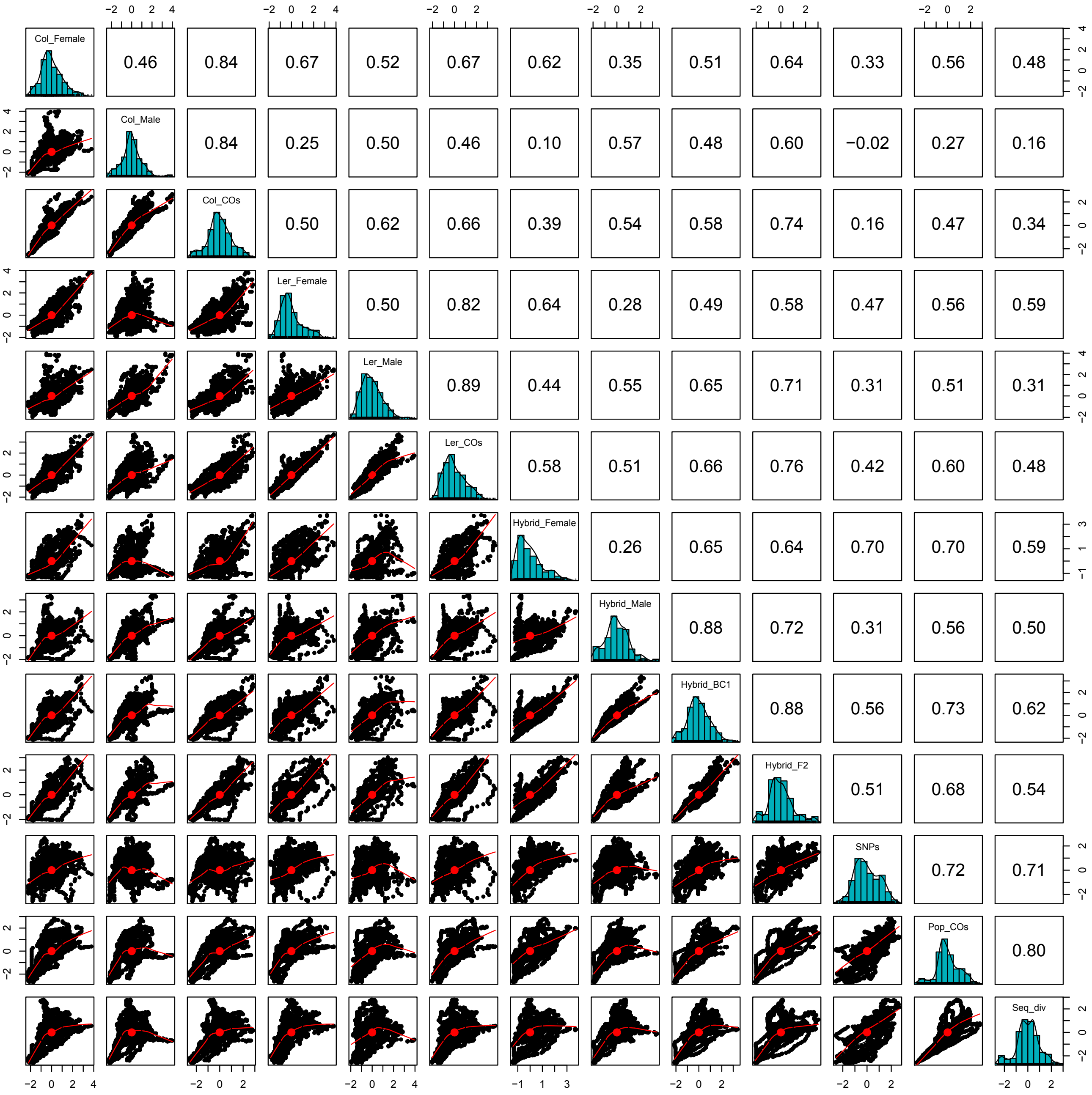


**Figure S14. Correlation analysis of CO distribution with polymorphisms.**
Spearman's correlation between each pair of features is shown in the upper triangle panel.

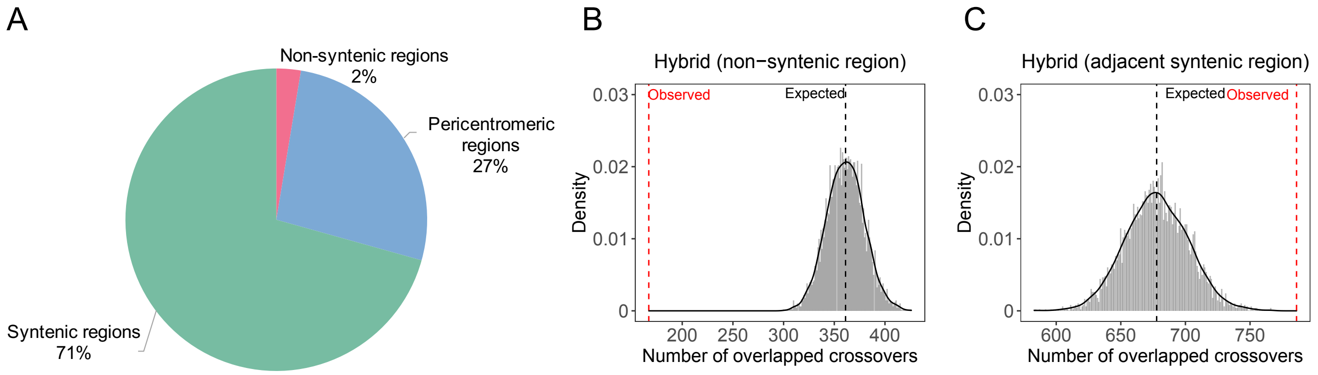


**Figure S15. Permutation analysis of the overlap between non-syntenic and adjacent syntenic regions and COs in Col/L*er* F2 hybrids.**

(A) The percentages of non-syntenic and syntenic regions in Col and L*er* and pericentromeric regions. (B-C) Permutation tests were performed to evaluate the overlap between non-syntenic and adjacent syntenic regions and COs in Col/L*er* F2 hybrids, respectively. The adjacent syntenic regions were defined as the up- and downstream regions (same size) of non-syntenic regions. The vertical black dashed lines indicate the expected mean number of overlaps in 5,000 shuffled permutations. The observed number of overlaps is represented by vertical red dashed lines.

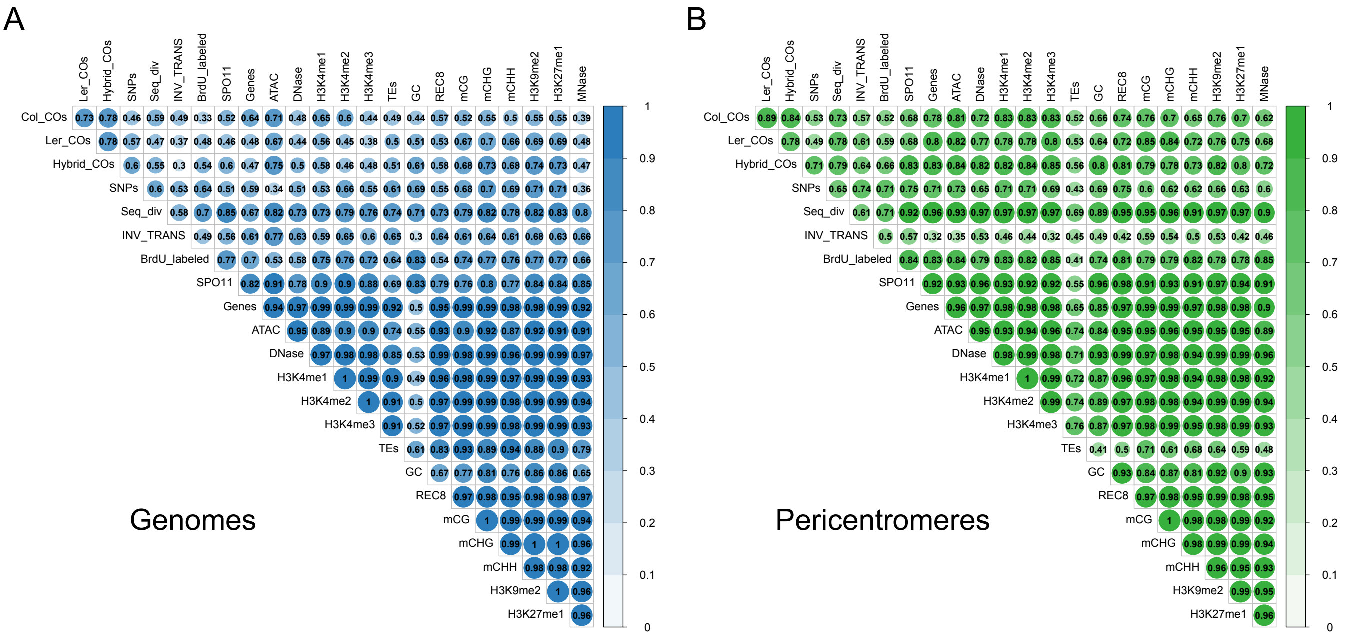


**Figure S16. Non-linear correlation coefficient matrices for the distribution of COs, genomic and epigenomic features at genome and pericentromeric scales.**
Col_COs, Ler_COs and Hybrid_COs (CO landscapes in Col, L*er*, and F2 hybrids), SNPs ( SNPs density between Col and L*er*), Seq_div (sequence diversity in the population of 2,029 Arabidopsis accessions), INV_TRANS (inversions and translocations between Col and L*er*), BrdU_labelled (origins of DNA replication), SPO11 (SPO11-1-oligos), Genes, TEs and GC (gene, TE and GC content density), ATAC and DNase (chromatin accessibility, ATAC-seq and Dnase-seq), H3K4me1/2/3, H3K9me2, H3K27me1 (euchromatin, heterochromatin and Polycomb histone marks, ChIP-seq), REC8 (cohesin, ChIP-seq), mCG, mCHG and mCHH (DNA methylation in CG, CHG and CHH contexts), MNase (nucleosome occupancy, MNase-seq).

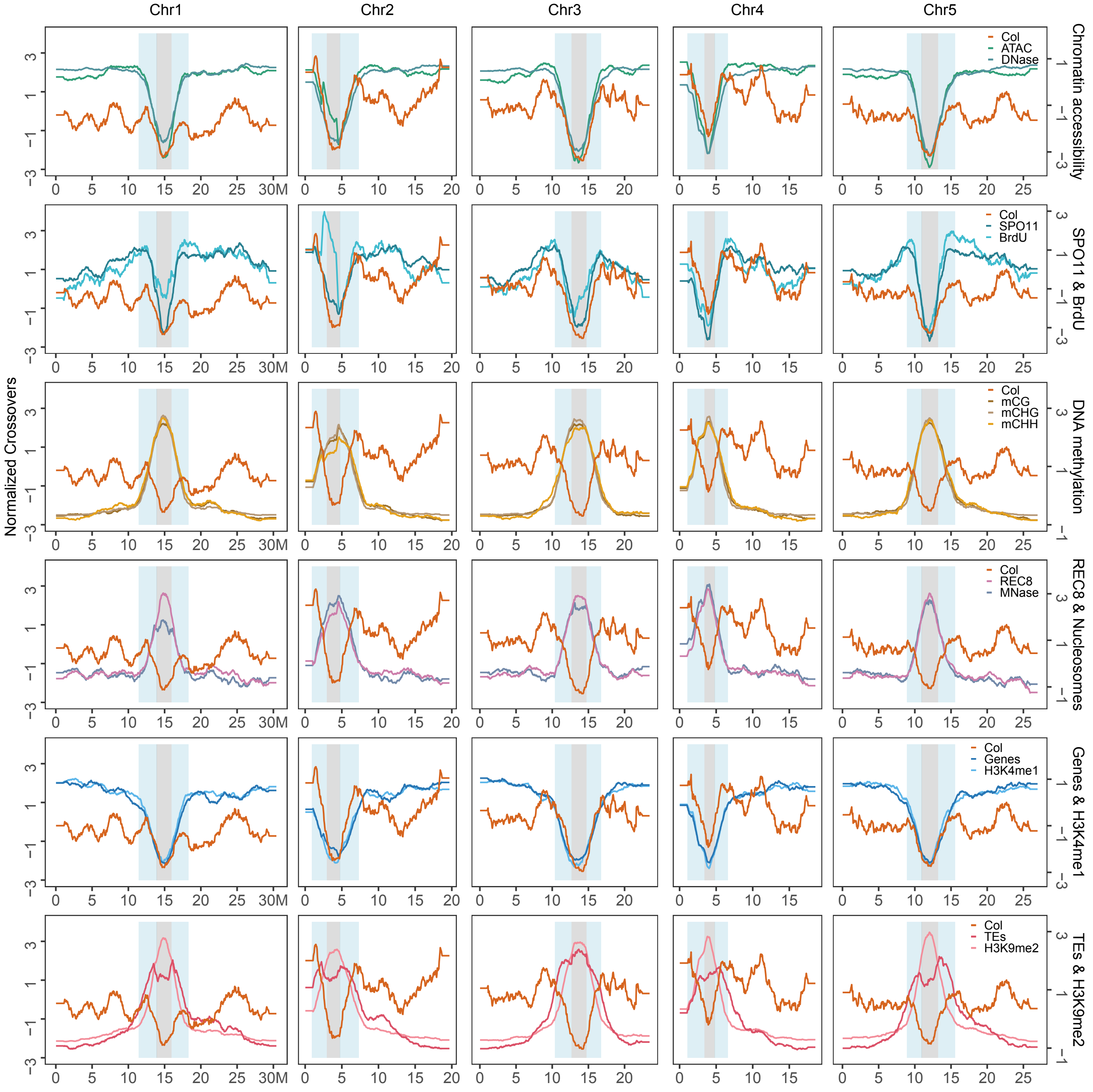


**Figure S17. Genomic landscape of Col COs, genomic and epigenomic features.**
Sliding window-based distributions (window size 2 Mb, step size 50 kb) were normalized to the same scale. The pericentromeric and centromere regions are indicated by grey and blue shading separately. Col (CO landscapes in Col), BrdU (origins of DNA replication), SPO11 (SPO11-1-oligos), Genes and TEs (gene and TE density), ATAC and DNase (chromatin accessibility, ATAC-seq and Dnase-seq), H3K4me1 and H3K9me2 (euchromatin and heterochromatin histone marks, ChIP-seq), REC8 (cohesin, ChIP-seq), mCG, mCHG and mCHH (DNA methylation in CG, CHG and CHH contexts), MNase (nucleosome occupancy, MNase-seq).

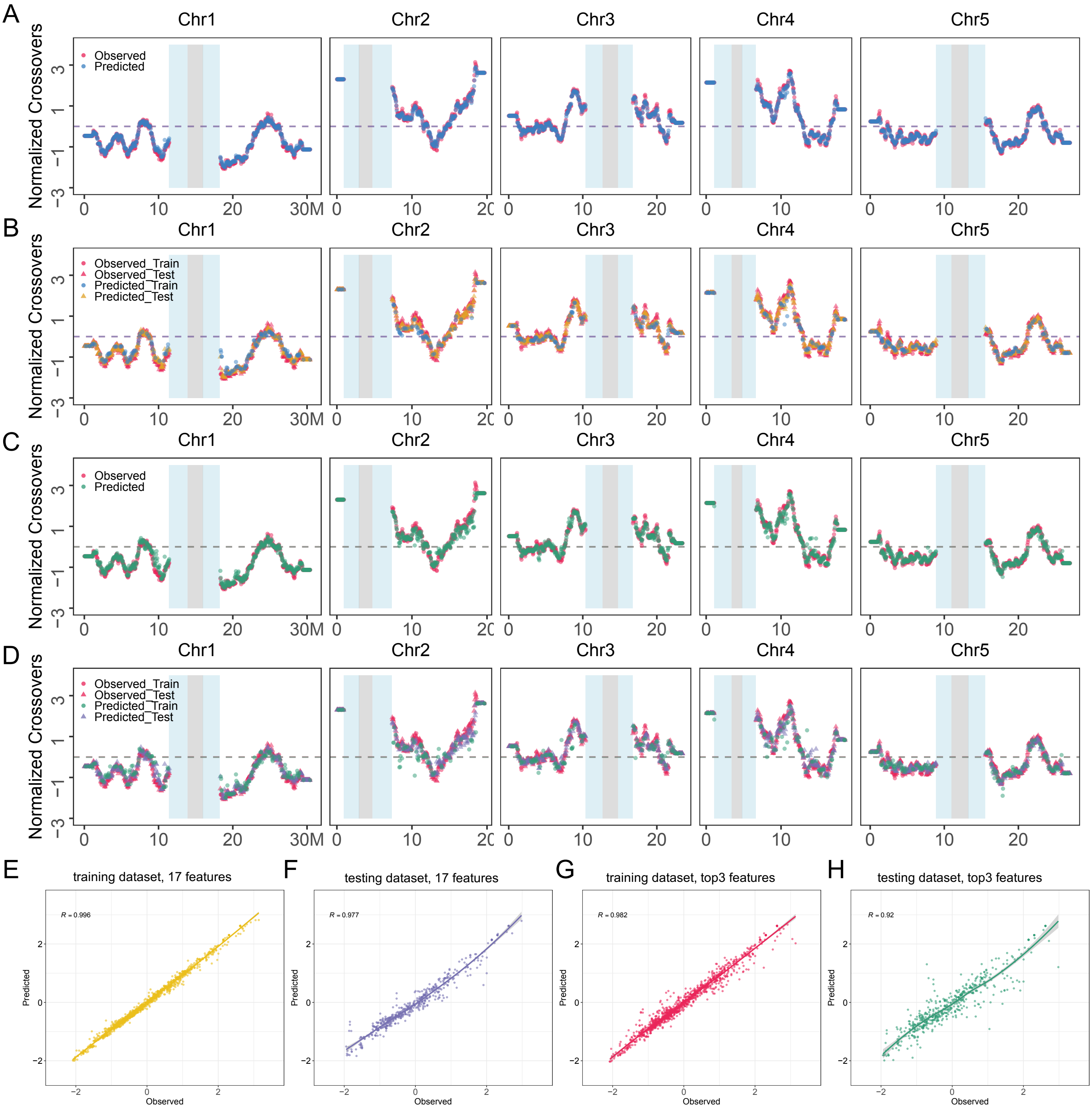


**Figure S18. The performance of the random forest model for predicting CO distribution in Col.** (A-B) The distribution of observed and predicted CO frequency by modelling with 17 features in the whole, training (70%) and testing (30%) datasets, respectively. (C-D) The distribution of observed and predicted CO frequency by modelling with the top three most important features in the whole, training and testing datasets, respectively. The pericentromeric and centromere regions are indicated by grey and blue shading separately. (E-H) Spearman's correlation between the predicted and observed CO distribution predicted by models constructed with 17 features and the top three most important features in the training and testing datasets, respectively.

**Table S1. Summary of EMS-induced mutations in Arabidopsis inbred lines**

| **Arabidopsis accession** | **F1*** | **Sequencing depths** | **Num. of variations after filtering** | **Num. of phased mutations** | **% of phased mutations** | **Num. of phased mutations per F1** |
|---|---|---|---|---|---|---|
| Col | 4314_A | 65.72 | 838 | 838 | 100 | 391 |
| Col | 4314_B | 44 | 955 | 955 | 100 | 448 |
| Col | 4314_E | 64.17 | 982 | 982 | 100 | 460 |
| Col | 4314_F | 62.27 | 953 | 953 | 100 | 449 |
| L*er* | 4560_B | 16.88 | 512 | 509 | 99.41 | 239 |
| L*er* | 4560_C | 16.22 | 471 | 471 | 100 | 214 |
| L*er* | 4560_D | 13.05 | 539 | 536 | 99.44 | 252 |
| L*er* | 4560_E | 17.03 | 526 | 521 | 99.05 | 243 |

**Table S2. The public dataset used in this study**

| Data set | Arabidopsis accession | ArrayExpress accession | GEO accession | SRA accession | Tissue | Reference |
|---|---|---|---|---|---|---|
| REC8 ChIP-seq | Col-0 | E-MTAB-7370 | | ERR3813870, ERR3813871 | Floral buds | (Lambing, et al. 2020) |
| REC8 ChIP-seq input | Col-0 | E-MTAB-7370 | | ERR3813873 | Floral buds | (Lambing, et al. 2020) |
| H3K4me3 ChIP-seq | Col-0 | E-MTAB-5048 | | ERR1590145, ERR1590146, ERR1590147 | Floral buds | (Choi, et al. 2018) |
| H3K4me3 ChIP-seq input | Col-0 | E-MTAB-6257 | | ERR2215860, ERR2215861, ERR2215862, ERR2215863 | Floral buds | (Choi, et al. 2018) |
| H3K9me2 ChIP-seq | Col-0 | E-MTAB-7370 | | ERR3813867 | Floral buds | (Lambing, et al. 2020) |
| H3K9me2 ChIP-seq input | Col-0 | E-MTAB-7370 | | ERR3813868 | Floral buds | (Lambing, et al. 2020) |
| H3K4me1 ChIP-seq | Col-0 | E-MTAB-7370 | | ERR3813865 | Floral buds | (Lambing, et al. 2020) |
| H3K4me2 ChIP-seq | Col-0 | E-MTAB-7370 | | ERR3813866 | Floral buds | (Lambing, et al. 2020) |
| H3K27me1 ChIP-seq | Col-0 | E-MTAB-7370 | | ERR3813864 | Floral buds | (Lambing, et al. 2020) |
| Histone ChIP-seq input | Col-0 | E-MTAB-7370 | | ERR3813869 | Floral buds | (Lambing, et al. 2020) |
| MNase-seq | Col-0 | E-MTAB-5042 | | ERR1590154 | Floral buds | (Choi, et al. 2018) |
| MNase-seq control | Col-0 | E-MTAB-6257 | | ERR2215860, ERR2215861, ERR2215862, ERR2215863 | Floral buds | (Choi, et al. 2018) |

| | | | | | | |
|---|---|---|---|---|---|---|
| DNase-seq | Col-0 | | GSE34318 | SRR388660, SRR388661 | Flowers | (Zhang, et al. 2012) |
| DNase-seq control | Col-0 | | | SRR10051102, SRR10051103 | Roots | (Alvarez, et al. 2019) |
| ATAC-seq | Col-0 | | GSE155503 | SRR12362020, SRR12362021, SRR12362022, SRR12362023 | Flowers | (Zhong, et al. 2021) |
| ATAC-seq control | Col-0 | | GSM2704269 | SRR5829244 | Roots | (Maher, et al. 2018) |
| DNA mehtylation BS-seq | Col-0 | | GSM2306321, GSM2306322 | SRX2148720, SRX2148721 | Meiocytes | (Walker, et al. 2018) |
| SPO11-1-oligo | Col-0 | E-MTAB-5041 | | ERR1590157, ERR1590158, ERR1590166 | Floral buds | (Choi, et al. 2018) |
| SPO11-1-oligo control | Col-0 | E-MTAB-6257 | | ERR2215864 | Floral buds | (Choi, et al. 2018) |
| Origins of replication BrdU-seq | Col-0 | | GSM542644 | SRR051937 | MM2d cells | (Costas, et al. 2011) |
| BrdU-seq control | Col-0 | | GSM542645 | SRR051938 | MM2d cells | (Costas, et al. 2011) |
| BrdU-seq control | Col-0 | | GSM588603 | SRR068545 | MM2d cells | (Costas, et al. 2011) |
